# State-selective nanobodies to probe Guanylate-binding protein coat assembly and function

**DOI:** 10.64898/2026.09.30.755308

**Authors:** Lennart L. Pagani, Tanja I. Kuhm, Cecilia Pinto, Elise G. Perton, Mireya Arents, Phuong N. M. Nguyen, Jan Steyaert, Els Pardon, Arjen J. Jakobi

## Abstract

Guanylate-binding proteins (GBPs) are interferon-inducible guanosine triphosphate hydrolases that coat cytosolic bacteria and promote non-canonical inflammasome activation. GBP coat assembly is initiated by GBP1, which binds bacterial lipopolysaccharide and recruits further paralogs in a hierarchical, nucleotide-dependent manner. Dissecting individual GBP functions remains challenging because the seven human paralogs are structurally similar and undergo nucleotide-driven conformational changes. Here, we developed a nanobody toolbox for state-selective probing of human GBP coat assembly. We identify eight nanobodies with distinct paralog, domain, and conformational specificities and define their binding profiles and epitopes using biophysical assays and cryogenic electron microscopy. Selected nanobodies were functionally validated in membrane reconstitution assays and infected cells. An inhibitory nanobody blocks progression from soluble GBP1 assemblies to LPS-associated coats, suggesting a stage-specific structural transition during coat formation. In contrast, a dimer-selective nanobody binds assembled coats without measurably perturbing their organisation and is recruited to GBP1-positive bacteria, indicating that this dimer interface remains intact and accessible in coat-associated GBP1. Together, these nanobodies provide state-selective reagents to distinguish, perturb, and probe GBP assemblies in structural and cellular studies.

## Introduction

Guanylate-binding proteins (GBPs) are important regulators of cell-autonomous immunity and contribute to host defence against diverse intracellular pathogens, including bacteria, parasites and viruses ^1–3^. GBPs are cytosolic guanosine triphosphate hydrolases (GTPases) of the dynamin superfamily and are broadly expressed in vertebrate cells ^1,4^. In response to intracellular infection, interferon signaling induces the expression of GBPs^5,6^, enabling them to assemble on pathogen surfaces or pathogen containing vacuoles (PCVs) ^7–11^. These assemblies can promote antimicrobial pathways such as non-canonical inflammasome activation and pyroptotic cell death ^12,13^.

In humans, seven GBP paralogs (GBP1-7) have been identified that can be recruited to the pathogen or PCV in a hierarchical manner ^4,10,11,14^. For cytosol-dwelling gramnegative bacterial pathogens, GBP1 has been shown to initiate encapsulation by associating with the bacterial outer membrane and recruiting other GBP paralogs, thereby forming large and stoichiometrically complex oligomeric assemblies ^10,11,14–16^. Several GBP paralogs contribute to non-canonical inflammasome activation by organising lipopolysaccharide (LPS)-containing bacterial surfaces or cytosolic LPS into caspase-4-activating platforms ^9,10,17,18^. Recent work further shows that GBP1 can remodel LPS-rich membranes in a manner that promotes caspase-4 binding and activation ^19,20^. Despite these advances, the molecular organisation of GBP assemblies and the contributions of individual GBP paralogs and oligomeric states remain incompletely understood. At the molecular level, GBPs share a high degree of sequence and structural similarity and can form homo- and heterodimers in a nucleotide-dependent manner ^14,21^. The Large GTPase (LG) domain mediates nucleotide binding and drives dimerisation during GTP hydrolysis ^22–24^. Dimerisation rearranges GBP1 into an outstretched conformation, with the middle domains (MDs) forming a parallel crossover beneath the LG dimer and the GTPase effector domains (GEDs) positioned for membrane engagement ^19,25^. In GBP1, GBP2 and GBP5, the GED harbours a covalently attached lipid moiety at its carboxyl terminus, which becomes exposed upon the conformational change associated with nucleotide hydrolysis and enables membrane anchoring ^19,26,27^. Thus, nucleotide hydrolysis functions as a molecular switch that couples oligomerisation, conformational rearrangement and membrane binding, a process that has been characterised most extensively for GBP1 (Fig. 1).

**Fig. 1.**
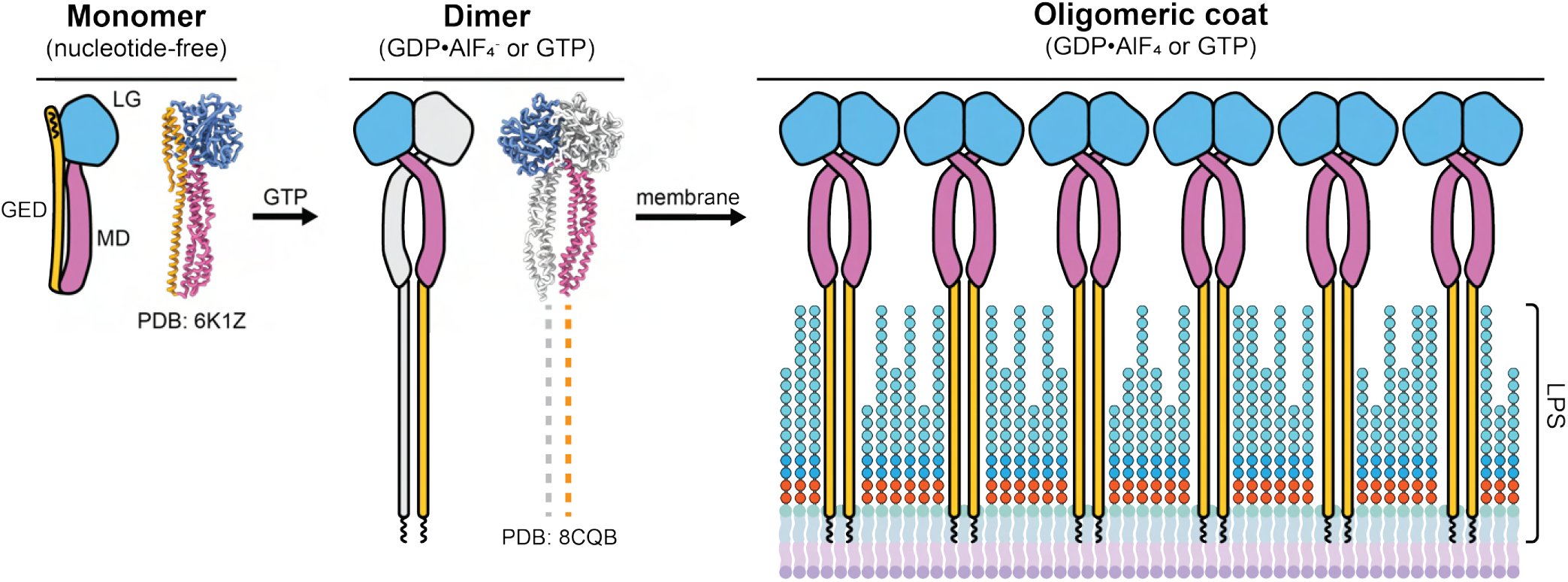
Nucleotide-dependent oligomeric states of GBP1. Left: Atomic model (PDB:6K1Z) and schematic representation of monomeric GBP1 in the absence of nucleotide. Middle: Upon GTP binding and hydrolysis, or in complex with the transition state mimic GDP *·* AlF_4_^−^, GBP1 dimerises to form an extended conformation with outstretched GEDs (PDB:8CQB). Right: In the presence of membranes, nucleotide-activated GBP1 further oligomerises on the membrane surface to form higher-order membrane-associated assemblies. LPS=lipopolysaccharide.

Despite recent progress, the molecular functions of individual GBP paralogs, their oligomeric states and their contributions to heterogeneous GBP coats remain incompletely understood ^18,28^. GBPs are challenging to study because their antimicrobial functions depend on both intrinsic properties, such as nucleotide-dependent conformational switching and oligomerisation, and extrinsic cues from the pathogen and cellular environment. Reconstitution in purified systems provides precise control of experimental conditions but often lacks the native membrane context and regulatory factors present in cells, whereas cellular experiments make it difficult to isolate the contribution of individual GBP paralogs and their conformational and oligomeric states. Therefore, probes that selectively visualise, perturb, or stabilise defined GBP states would be valuable tools for dissecting GBP-mediated immunity.

Single-domain antibodies, or nanobodies, are well suited to address this need because they combine high target specificity with broad applicability as structural and functional probes ^29^. Nanobodies derive from the variable domain of camelid heavy-chain antibodies and adopt an immunoglobulin fold ^29,30^. Their single-domain architecture enables targeting of epitopes that may be inaccessible to conventional antibodies and permits intracellular expression ^30,31^. Target recognition is mediated by three variable complementarity-determining regions (CDRs) that support high-affinity and specific binding. These properties have established nanobodies as versatile tools in cell and structural biology ^32–35^, including in modular systems for localisation, visualisation, and spatial or temporal manipulation in cells ^36,37^.

Here, we report a set of nanobody-based probes for the structural and functional investigation of human GBPs, centred on GBP1. We characterise eight nanobodies that differ in paralog and conformation specificity, binding kinetics, and their effects on oligomerisation of GBP1, GBP2 and GBP3. Using complementary biophysical assays and cryo-EM, we define their binding properties and map the epitopes of selected nanobodies. We further illustrate their functional utility through two nanobodies with contrasting effects on GBP coats, with one perturbing GBP1 coat formation and the other recognising assembled GBP1 coats without measurably altering their organisation, and validate that both also function as intracellular probes in infected cells. Together, this toolbox provides versatile set of reagents for probing GBP assemblies across biochemical, structural, and cellular contexts.

## Results

### Generation of nanobodies targeting distinct GBP conformational and oligomeric states

To generate nanobodies that recognise distinct GBP conformations and epitopes, we immunised llamas (*Lama glama*) with multiple human GBP1 variants and domains. These included farnesylated and non-farnesylated fulllength GBP1 in its inactive (apo-) form or bound to GDP *·* AlF_4_ ^−^, which stabilises GBP1 dimers by mimicking the transition state of GTP hydrolysis (Fig. 1). In addition, we used the isolated large GTPase domain (GBP1-LG), either in the nucleotide-free monomeric or dimeric GDP *·* AlF_4_ ^−^-bound state, to selectively identify nanobodies that recognise the LG domain, with the rationale that these are less likely to sterically hinder and interfere with GBP1 coat assembly ^19,25^. Following consecutive rounds of immunisation, lymphocytes were isolated from blood and the RNA was extracted to synthesise cDNA. The resulting cDNA was used to create phage display libraries, which were enriched by biopanning against the respective GBP1 antigens. Individual clones were identified by enzyme-linked immunosorbent assay screening and grouped into nanobody families based on complementarity-determining region 3 (CDR3) sequence conservation. In total, we identified 114 VHH sequences that were grouped into 74 CDR3 sequence families, comprising 15 multi-member families and 59 singleton families (Fig. S1). We purified representative nanobodies for each of these families and first assessed their binding capabilities towards GBP1 by size-exclusion chromatography coupled to multi-angle light scattering (SEC-MALS; Fig. 2a, Fig. S1). To obtain a functionally useful and sequence-diverse set of probes, we selected eight nanobodies for further characterization and examined their binding to monomeric and dimeric GBP1, GBP2, GBP3ΔGED and GBP1-LG by SEC-MALS and mass photometry (Fig. 2a,b, Fig. S2-S4). In parallel, we determined their binding affinities and kinetics toward monomeric and dimeric GBP1 and GBP2 by surface plasmon resonance (SPR) using single cycle kinetics (Fig. 2c, Fig. S5, Table S1). Together, these complementary biophysical assays allowed us to define the binding profiles of the selected nanobodies, which are summarised in Figure 2d and discussed in the following.

**Fig. 2.**
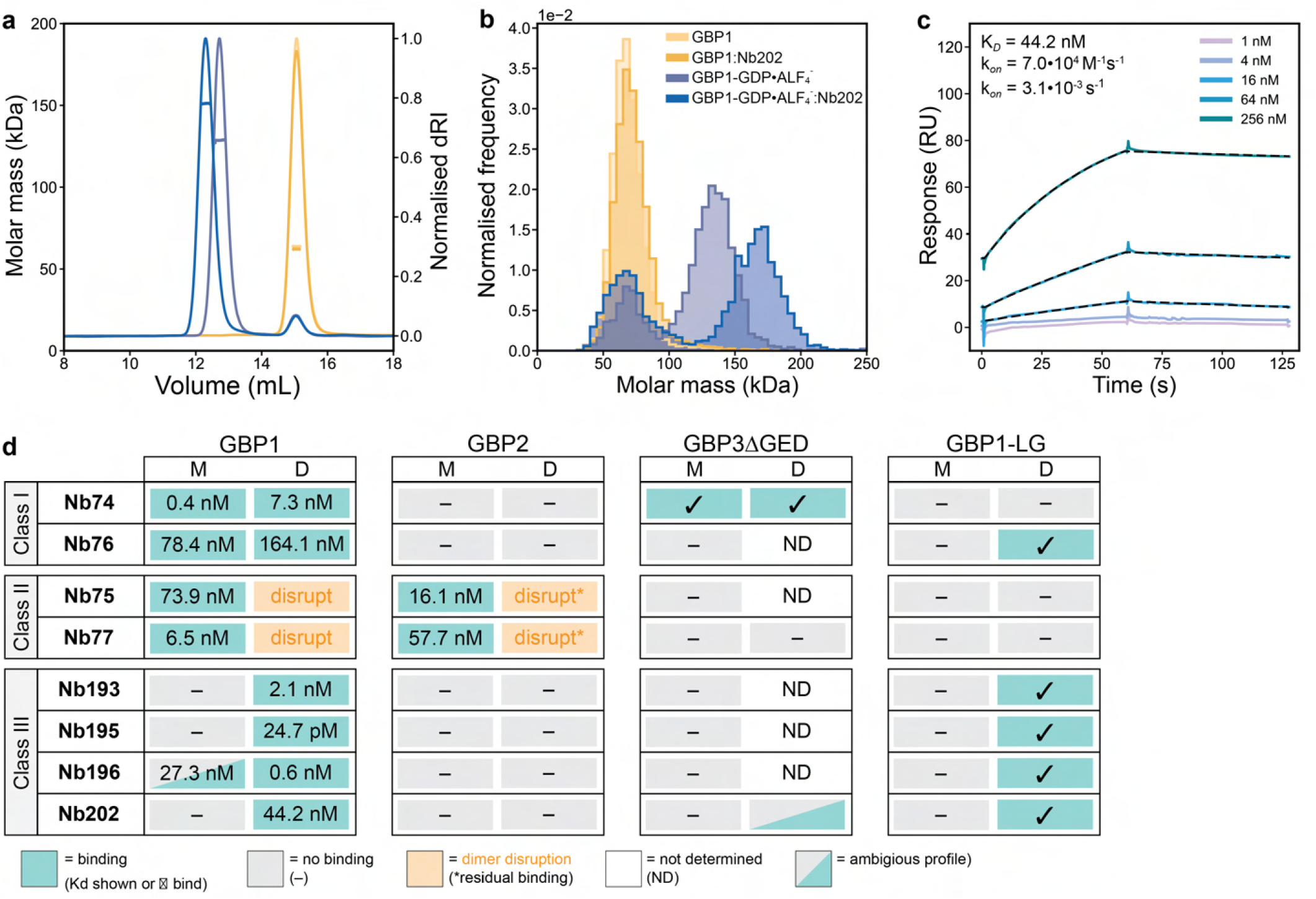
Biophysical characterisation of GBP-binding nanobodies. **a-c**, Representative data from the biophysical assays used to characterise nanobody binding to GBP paralogs and domains in monomeric and GDP *·* AlF_4_^−^-stabilised dimeric states, Nb202 is shown as example. **a**, SEC-MALS data showing a shift in the elution profile and molar mass of dimeric GBP1 upon nanobody binding. **b**, Mass photometry data showing changes in measured molar mass for dimeric GBP1 upon nanobody binding. **c**, Representative SPR sensorgrams for Nb202 binding to immobilised GBP1 GDP *·* AlF_4_ ^−^ showing the concentration-dependent kinetic analysis. Data were acquired using single-cycle kinetic titrations and are plotted as multi-cycle kinetic sensorgrams for brevity. Kinetic parameters are shown inset. **d**, Consensus binding matrix summarising nanobody binding profiles and parameters across GBP variants and nucleotide states (M: nucleotide-free monomer, D: GDP *·* AlF_4_ ^−^-activated dimer). The matrix integrates data from SEC-MALS, mass photometry, and SPR (see Figs. S3-S4). Nanobodies are grouped by classes representing their binding profiles. Class I nanobodies bind both monomeric and dimeric GBP1. Class II nanobodies disrupt and prevent GBP1 and GBP2 dimerisation. Class III specifically recognises dimeric GBP1.

The selected nanobodies were separated into three distinct binding classes, reflecting differences in paralog specificity, conformational preference, affinity, and effects on GBP dimerisation. The first class comprising of Nb74 and Nb76 (class I) recognised both monomeric and GDP *·* AlF_4_ ^−^-induced dimeric GBP1. Nb74 bound with sub-nanomolar affinity to monomeric GBP1 (K_D_=0.4 nM) and also recognised GBP3ΔGED, but not GBP2, suggesting that its epitope is conserved between GBP1 and GBP3 but diverges in GBP2. This binding profile is rationalised by structural epitope mapping in the following section. Nb76 also recognised both GBP1 states, albeit with substantially lower affinities of 78.4 nM and 164.1 nM for monomeric and dimeric GBP1, respectively.

The second class comprised Nb75 and Nb77 (class II). Although these nanobodies belong to different CDR3 sequence families, both exclusively bound monomeric GBP1 and disrupted GDP *·* AlF_4_ ^−^-stabilised GBP1 dimers. Nb75 and Nb77 bound monomeric GBP1 with affinities of 73.9 nM and 6.5 nM, respectively, and also recognised GBP2 with the same state-dependent binding profiles although their disruptive effect on GDP *·* AlF_4_ ^−^-stabilised GBP2 dimers appeared weaker than for GBP1. Neither nanobody bound the isolated GBP1-LG domain, suggesting that their epitopes are located within the MD and GED, or a conformation-dependent interface with the LG domain involving these regions. Because both nanobodies disrupt preformed GBP1 and GBP2 dimers, they provide candidate perturbation tools to interfere with GBP oligomerisation and coat assembly.

A third class including Nb193, Nb195, Nb196 and Nb202 (class III) preferentially recognised dimeric GBP1. Nb202 selectively bound dimeric GBP1 with nanomolar affinity (K_D_=44.2 nM) and also bound dimeric GBP1-LG, but showed no detectable binding to monomeric GBP1, monomeric/dimeric GBP2, or GBP3ΔGED under the conditions tested. This suggests that subtle sequence differences between GBP1 and GBP2/GBP3ΔGED can alter conformational selectivity. Nb193 and Nb195 also selectively recognised dimeric GBP1 at the LG domain and bound with high affinity (K_D_=2.1 nM and 24.7 pM, respectively). These nanobodies therefore provide complementary probes for detecting or stabilising dimeric GBP1.

Nb196 showed an intermediate binding profile. SEC-MALS and mass photometry indicated preferential binding to dimeric GBP1, whereas SPR showed sub-nanomolar affinity (K_D_=0.6 nM) to dimeric GBP1 and transient binding to monomeric GBP1 with a K_D_ of 27.3 nM and a high dissociation rate (Fig. S5, Table S1). This suggests that Nb196 may dissociate from monomeric GBP1 during SEC-MALS and mass photometry measurements, while binding to dimeric GBP1 is sufficiently stable to be detected across assays. Nb196 also appeared to bind GBP1 dimers with a 1:2 nanobody:GBP1 stoichiometry, a feature that we examine structurally below.

### Structural mapping of nanobody binding epitopes

To understand the molecular basis of the distinct binding profiles observed in our biophysical assays, we used cryo-EM to map the epitopes of selected GBP-binding nanobodies. We focused on nanobodies that recognise dimeric GBP1, as their complexes with GDP *·* AlF_4_ ^−^-stabilised GBP1 were expected to be most suitable for single-particle analysis. We obtained cryo-EM structures for Nb193, Nb195, Nb196, and Nb202 bound to dimeric GBP1, which we compared with the previously determined Nb74–GBP1 complex ^19^ (Fig. 3a-f, Fig. S6, Table S2). The Nb202 complex was resolved to 4.1 Å, allowing detailed interpretation of the Nanobody–GBP1 interface (Fig. 3a,f, Fig. S7a, Table S3). Although the maps for Nb193, Nb195, and Nb196 were lower in resolution, they supported rigidbody fitting of AlphaFold2-predicted nanobody models and thereby define the approximate locations and orientations of their epitopes (Fig. 3a, c-e, S7c). We could not resolve structures of Nb75 or Nb77because these nanobodies bind and stabilise monomeric GBP1, resulting in complexes that were less amenable to single-particle analysis..

**Fig. 3.**
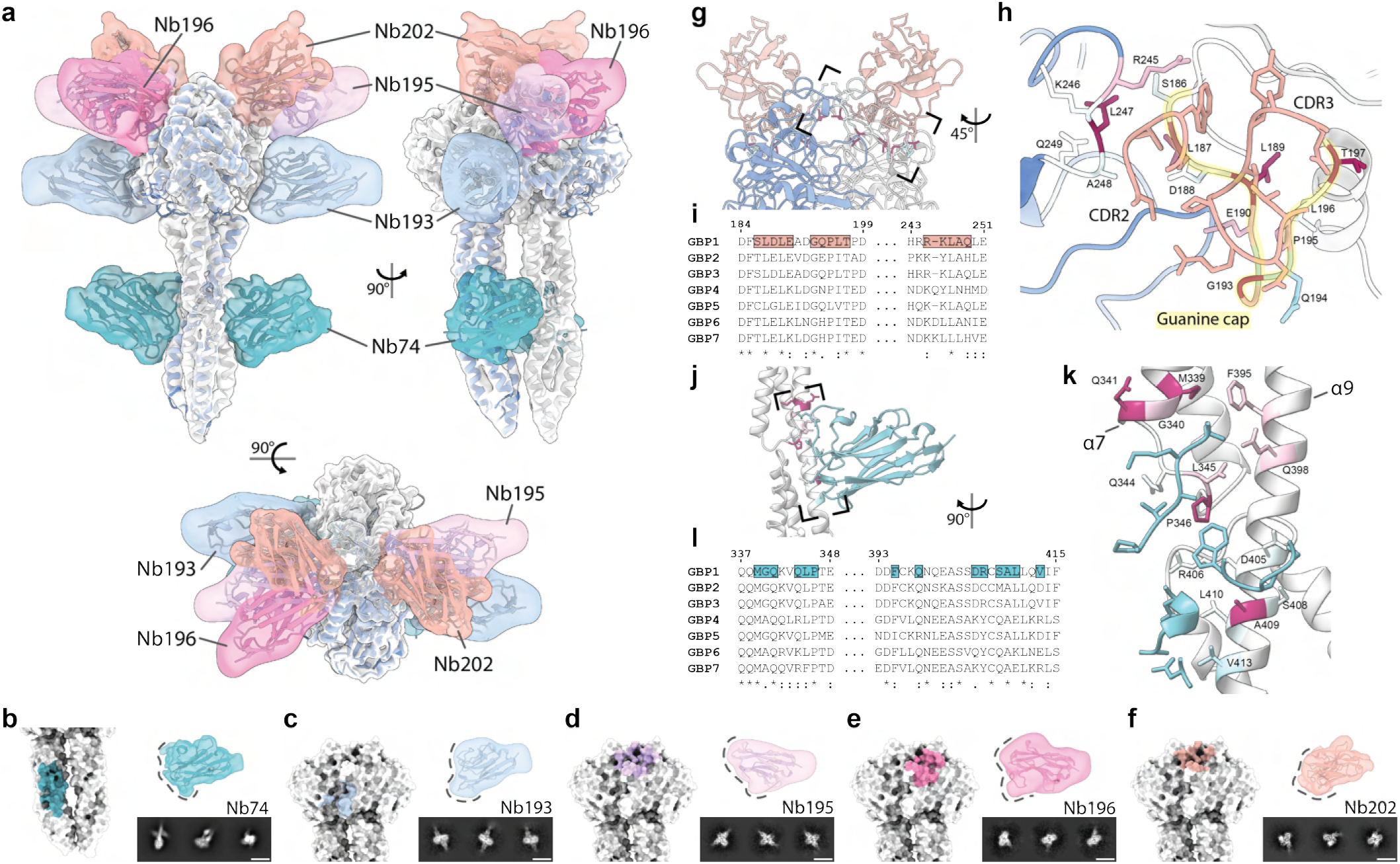
Structural analysis of GBP-binding nanobodies. **a**, Cryo-EM ensemble map of selected nanobodies in complex with dimeric GBP1. Cryo-EM density maps of Nb193 (EMDB:59826; blue), Nb195 (EMDB:59827; purple), Nb196 (EMDB:59828; pink) and Nb202 (EMDB:59825; orange) superimposed on GBP1 (grey) in complex with Nb74 (EMDB:16794; green). The individual monomer chains of the GBP1 model are coloured in grey and blue. **b-f**, Cryo-EM density maps of individual nanobodies (right) and their binding interfaces superposed on the surface model of GBP1 GDP *·* AlF_4_ ^−^ (left). Representative 2D class averages are shown below. Scale bar is 10 nm. **g**, Atomic model of Nb202 bound to its epitopes on GBP1-GDP *·* AlF_4_ ^−^ dimers (PDB:000034GI). **h**, Close-up of the box-out-region in (g) showing how Nb202 contacts the guanine cap (highlighted in yellow) at the GBP1 dimer interface. The GBP1 model is coloured according to the sequence conservation. **i**, Multiple sequence alignments of human GBP paralogs with Nb202-interacting residues highlighted. **j**, Atomic model of Nb74 bound to its epitopes on GBP1-GDP *·*AlF_4_ ^−^ dimers (PDB:000034GJ). **k**, Close-up of the box-out-region in (j) showing Nb74 binding at the groove formed by α7/α9 in the GBP1 middle domain. The GBP1 model is coloured according to the sequence conservation. **l**, Multiple sequence alignments of human GBP paralogs with Nb74-interacting residues highlighted.

The structural analysis revealed a prominent cluster of nanobody epitopes near the LG domain dimer interface. Nb193, Nb195, Nb196, and Nb202 all bind close to this region, consistent with their preferential recognition of dimeric GBP1. Nb193, Nb195, and Nb202 appear to contact residues from both GBP1 chains, providing a structural explanation for their dimer-selective binding (Fig. 3a). By contrast, Nb196 binds primarily to one GBP1 chain. This binding mode is consistent with its weak, transient interaction with monomeric GBP1 observed by SPR. The presence of only one Nb196 density per GBP1 dimer in the cryo-EM structure further agrees with the apparent 1:2 nanobody:GBP1 stoichiometry observed by SEC-MALS and mass photometry. Although the GBP1 LG dimer is pseudo-symmetric ^19,23^, the second equivalent site was unoccupied, suggesting that Nb196 binding may (allo)sterically prevent binding of a second nanobody.

The different positions and orientations of the LG-domain binders suggest distinct potential effects on GBP1 assembly. Nb193 and Nb195 bind laterally to the GBP1 dimer and may therefore interfere with higher-order oligomerisation of GBP1 dimers^19,25^ without directly disrupting the dimer interface. In contrast, Nb202 binds in an upright orientation on the LG domain, leaving the lateral surfaces of the dimer comparatively unobstructed. The higher-resolution Nb202 map further revealed that Nb202 recognises a composite epitope at the GBP1 dimer interface. Its CDRs contact the guanine cap, a flexible loop characteristic of GBPs, and additionally engages residues on the opposing GBP1 chain (Fig. 3g-i). This explains why Nb202 selectively recognises dimeric GBP1. Interestingly, the closely related Nb76 differs from Nb202 by only two residues but binds both monomeric and dimeric GBP1 albeit with lower affinity (Fig. S1, Fig. S2a). These residue variants, N73H in framework region 3 and A101P in CDR3, are positioned at the Nanobody– GBP1 interface and may alter the flexibility or contact geometry of the paratope, thereby broadening recognition to monomeric GBP1. Sequence conservation across human GBP paralogs suggests that the Nb202 epitope is partially conserved in GBP3 and GBP5, although this requires experimental validation.

Nb74 binds a distinct epitope outside the LG-domain dimer interface located in the middle domain, in a groove formed by helices α7 and α9 (Fig. 3j-l, Fig. S7b). This region is not substantially rearranged between monomeric and dimeric GBP1, explaining why Nb74 recognises both conformational states. The epitope is also conserved between GBP1 and GBP3 but diverges in other GBP paralogs, consistent with the observed binding of Nb74 to GBP1 and GBP3ΔGED, but not GBP2. Thus, Nb74 provides a conformationally permissive GBP1/GBP3-directed binder, whereas the LG-domain binders preferentially report on the dimeric state of GBP1.

Nb75 and Nb77 represent a structurally distinct class of nanobodies that preferentially recognise monomeric GBP1. As these complexes could not be resolved by cryo-EM, we used structure prediction to investigate their potential binding modes. The predicted GBP1–Nb75 and GBP1–Nb77 complexes showed high model confidence, with interface predicted template modeling scores (ipTM) scores of 0.93 and 0.95, respectively (Fig. S7d,e). Consistent with the absence of binding to the isolated GBP1-LG domain, the predicted epitopes are located outside the LG domain and suggest that they may interfere with the conformational rearrangements of the helical domains that accompany formation of the outstretched GBP1 dimer. In the predicted complex, Nb77 binds near the α6-α7 cross-over loop, which connects the LG domain to the middle domain and is remodelled during the transition from the compact monomer to the outstretched dimeric conformation ^19^. Binding at this site could stabilise the monomeric arrangement of the helical domain or sterically hinder the displacement of the middle domain required for dimer formation. Nb75 is predicted to bind a groove formed between α9 of the MD and α12 of the GED, which may lock the GED in a conformation incompatible with membrane association. These predicted binding modes provide a structural rationale for their ability to disrupt GDP *·* AlF_4_ ^−^-stabilised GBP1 dimers. Their similar state-dependent recognition of GBP2 further suggests that the relevant structural features are conserved between GBP1 and GBP2^19,22,38^. Because these binding modes are based on computational predictions rather than experimentally determined structures, the precise epitopes and mechanisms of dimer disruption remain to be validated.Together, the structural analysis rationalises the distinct binding classes identified in the biophysical analysis and suggests how individual nanobodies may be used to interrogate GBP coat assembly. Inhibitory nanobodies such as Nb75 and Nb77 provide candidate tools to interfere with the early dimerisation steps required for coat formation. Other dimer-selective nanobodies recognise epitopes clustered near the LG dimer interface, with binding orientations that may either interfere with or preserve dense packing of GBP dimers in bacteria-associated coats.

### Nanobodies differentially affect GBP1 coat assembly

To test these structure-guided predictions experimentally, we selected two nanobodies with contrasting expected effects on GBP1 coat assembly. Nb77 was chosen as an inhibitory nanobody predicted to interfere with early assembly steps, whereas Nb202 was chosen as a dimer-selective nanobody whose upright binding mode appeared compatible with assembled coats. We first asked whether these nanobodies can bind to, or alter, preassembled GBP1 coats on artificial membranes.

To monitor nanobody binding to membrane-assembled GBP1 coats, we introduced a cysteine mutation at Ser^7^ in the nanobodies and Gln^577^ in GBP1, and site-specifically labelled Nb77 and Nb202 with CF488 and GBP1 with CF647. Nucleotide-activated GBP1^CF647^ was assembled on DHPE Texas Red-labelled giant unilamellar vesicles (GUVs), after which Nb77^CF488^ or Nb202^CF488^ were added to the preassembled coats. This setup allowed simultaneous time-lapse visualisation of the lipid membrane, GBP1 coat, and nanobodies by confocal microscopy (Fig. 4a,b, Fig. S8). To quantify nanobody recruitment and coat stability, we segmented individual GUVs using the Texas Red channel and measured the peak radial fluorescence signal at the membrane for each time point. The GBP1 and nanobody signals were normalised to the surrounding background to obtain a radial signal-to-noise ratio over time. This allowed us to assess both nanobody recruitment to GBP1-coated membranes and changes in membrane-associated GBP1 after nanobody addition. Nb77 did not colocalise with GBP1-coated GUVs and did not measurably reduce the GBP1 signal at the membrane. Thus, although Nb77 disrupts GBP1 dimers in solution, it does not detectably bind to or disassemble stable GBP1 coats once they have formed. In contrast, Nb202 robustly colocalised with GBP1-coated GUVs while leaving the GBP1 signal unchanged, indicating that Nb202 can access GBP1 within assembled coats without measurably altering coat-associated GBP1 density.

**Fig. 4.**
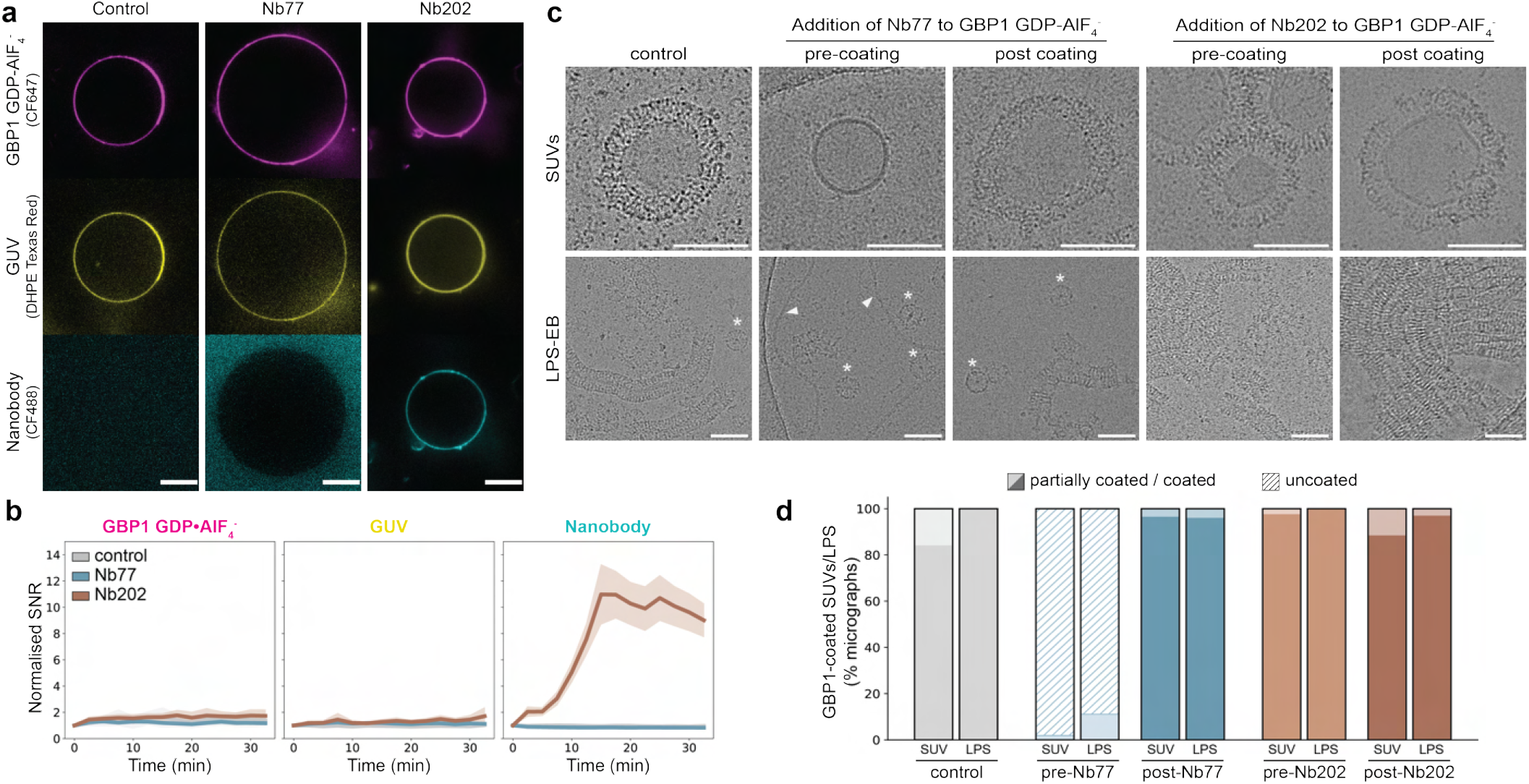
Nanobodies recognise GBP1 coats on artificial membranes and bacterial LPS. **a,b**, Nanobodies were added (t=0 min) to GUVs that were pre-coated with GBP1 GDP *·* AlF_4_ ^−^ and examined over time by confocal microscopy. **a**, Fluorescence images depicting GUVs (DHPE Texas Red) coated with GBP1 GDP *·* AlF_4_^−^ (CF647) and incubated with nanobodies (CF488). Scale bar is 5 µm. **b**, Corresponding graphs displaying the mean normalised radially averaged signal-to-noise ratio per GUV as a function of time after nanobody addition (Nb77: n=161, Nb202: n=133, Control: n=47). Shaded areas represent 95% confidence intervals. **c**, Cryo-EM micrographs of SUVs (top) and *E. coli* O111:B4 LPS (bottom, LPS-EB). Nanobodies were either pre-incubated with GBP1 prior to coat formation on SUVs/LPS-EB or added post coating. Micellar GBP1 assemblies and uncoated LPS micelles are annotated by asterisks and arrows, respectively. Scale bar is 100 nm. **d**, Corresponding bar graphs displaying the percentage of micrographs containing (partially) coated and uncoated SUVs/LPS-EB per condition (SUVs/LPS: control: n=56/26; pre-Nb77: n=52/18; post-Nb77: n=54/24; pre-Nb202: n=39/43; post-Nb202: n=51/31).

### Nanobodies reveal stage-specific transitions during GBP1 coat assembly

Because confocal microscopy reports on coat recruitment but not local coat architecture, we next used cryo-EM to assess whether the nanobodies affect GBP1 coats at higher spatial resolution. Nanobodies were added either before GBP1 coat formation or after coats had already formed on small unilamellar vesicles (SUVs) and smooth lipopolysaccharide purified from the pathogenic *E. coli* O111:B4 strain (LPS-EB; Fig. 4c,d).

When Nb77 was added to farnesylated GBP1 before coat assembly on SUVs or LPS-EB, extended GBP1 coats were not detected. Instead, micellar GBP1 assemblies remained visible, because they had formed via nucleotide-dependent oligomerisation before addition of Nb77^26,39^. This indicates that Nb77 does not readily disassemble this intermediate state, but prevents its further transition into extended SUV or LPS-associated coats. Addition of Nb77 after coat formation did not visibly disrupt preassembled coats, in agreement with the GUV experiments. Together with the ability of Nb77 to disrupt non-farnesylated GBP1 dimers in solution, these observations suggest that GBP1 becomes progressively resistant to Nb77 as assembly proceeds from soluble dimers through micellar intermediates to mature coats. On the contrary, Nb202 did not prevent coat formation when present before assembly and did not visibly alter preassembled coats when added afterwards. Thus, Nb202 is compatible with both the formation and maintenance of GBP1 coats. Together, these experiments show that the nanobodies distinguish different stages of GBP1 coat assembly and establish their utility for probing GBP coat assembly and function.

### Intrabodies probe GBP1 coat assembly in bacteria-in-fected cells

We next asked whether these contrasting behaviours are retained in the cellular context of bacterial infection, where GBP coat formation occurs in the presence of endogenous host factors, including additional GBP paralogs. We expressed the nanobodies intracellularly as GFP-tagged intrabodies in ΔGBP1 HeLa cells carrying doxycycline-inducible mCherry-GBP1. mCherry-GBP1 expression was induced with doxycycline in both uninfected and infected conditions. In uninfected cells, both GFP-Nb77 and GFP-Nb202 showed a diffuse intracellular distribution and did not accumulate in prominent GBP1-positive structures (Fig. S9). This indicates that the nanobodies do not form detectable intracellular assemblies under these conditions and provides a baseline for assessing their recruitment during infection.

We next infected cells with mCerulean3-expressing *Salmonella enterica* serovar Typhimurium and analysed nanobody localisation by confocal microscopy (Fig. 5, Fig. S10). GFP-Nb202 strongly colocalised with mCherry-GBP1-positive *S*. Typhimurium and formed continuous coat-like patterns around bacteria, consistent with its ability to bind GBP1 coats on artificial membranes. In some cases, GFP-Nb202 extended around bacteria beyond regions with detectable mCherry-GBP1 signal. Because Nb202 has not been tested systematically against all human GBP paralogs or mixed GBP assemblies, this additional signal cannot presently be assigned uniquely to GBP1 and may reflect recognition of another GBP paralog or coat regions in which mCherry-GBP1 lies below the detection threshold. In contrast, GFP-Nb77-expressing cells showed reduced mCherry-GBP1 coating of intracellular *S*. Typhimurium, consistent with the dimer-disrupting activity of Nb77 *in vitro*. Residual GFP-Nb77 signal was occasionally observed together with mCherry-GBP1 foci on bacteria, suggesting that Nb77 may associate with early or incomplete GBP1 assemblies rather than mature coats.

**Fig. 5.**
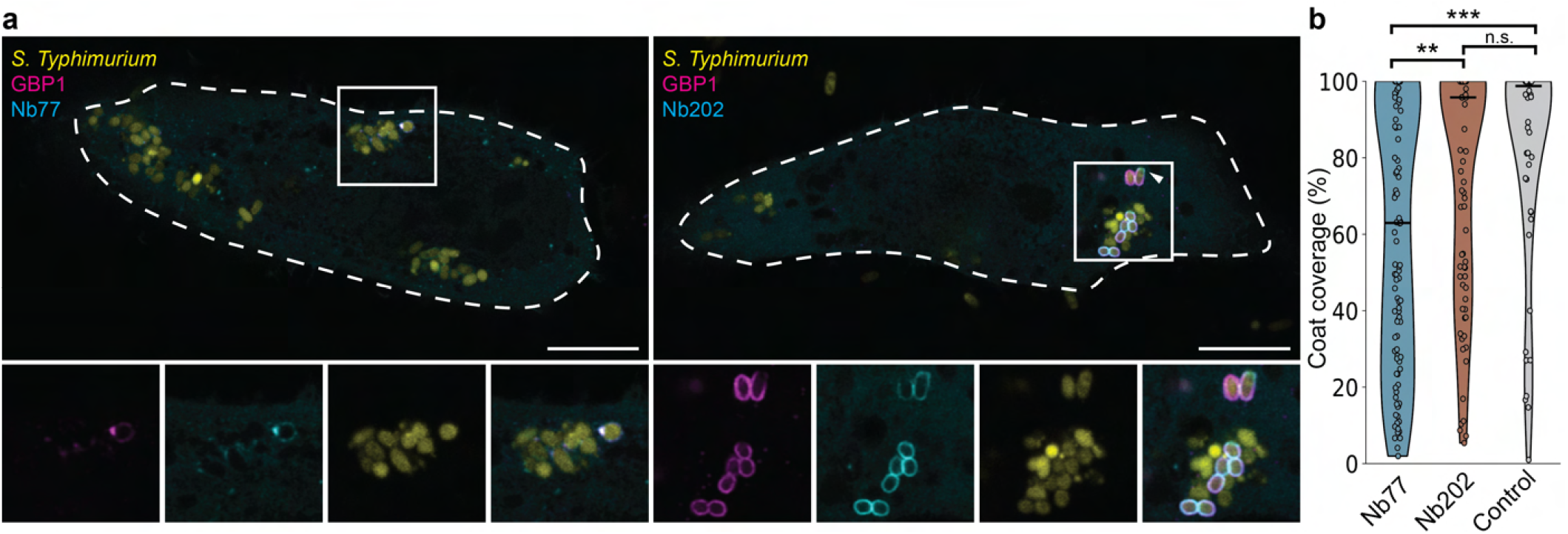
Intrabodies target GBP1 in *S*. Typhimurium infected cells. **a**, GFP-Nb77 (left) and GFP-Nb202 (right) were transfected in HeLa ΔGBP1 + Tet-mCherry-GBP1 cells. Cells were infected with *S*. Typhimurium-mCerulean3 and mCherry-GBP1 expression was induced via addition of doxycycline. Cells were fixated and imaged by confocal microscopy. The arrow indicates GFP-Nb202 putatively targeting other GBP paralogs. Lower panels show the individual fluorescence channels for the boxed region. Scale bar is 10 µm. **b**, Quantification of mCherry-GBP1 coat coverage on individual intracellular *S*. Typhimurium bacteria from the confocal images shown in (a). Each bacterium was individually traced, and coat coverage was calculated as the fraction of the bacterial circumference positive for mCherry-GBP1 signal. Data are shown as violin plots with individual bacteria plotted as points (Nb77: n=111, Nb202: n=82, Control: n=52). Statistical significance was assessed using a Kruskal-Wallis test followed by Holm-Bonferroni correction (Nb77-control: p=0.0004, Nb77-Nb202: 0.0027, Nb202-control: p=0.2832)

To quantify these effects, we traced individual bacteria and measured the bacterial circumference positive for mCherry-GBP1 signal (Fig. 5b). Cells expressing GFP-Nb77 displayed reduced mCherry-GBP1 coating of intracellular *S*. Typhimurium compared with control cells and cells expressing GFP-Nb202, consistent with the ability of Nb77 to disrupt or prevent GBP1 dimer formation and prevent coat assembly *in vitro*. By contrast, GFP-Nb202 did not significantly alter mCherry-GBP1 coat coverage compared to control cells. Together, these experiments provide a basis for future quantitative analysis of GBP coat formation and dynamics in infected cells.

## Discussion

The state-selective nanobodies developed here offer tools to probe conformational transitions during GBP coat assembly. Specifically, the activities of Nb77 and Nb202 reveal two distinct features of the assembly process that are difficult to infer otherwise. Nucleotide activation of farnesylated GBP1 promotes formation of micelle-like assemblies in the absence of membranes, while related outstretched GBP1 dimers form higher-order assemblies on lipid and LPS-containing membranes ^19,25,26^. Previous work proposed that polymeric GBP1 docks onto LPS-containing bacterial surfaces and subsequently depolymerises at the membrane, which delivers activated GBP1 molecules into the developing coat ^15^. Our observations suggest that such a transition would involve more extensive structural reorganisation than a direct transfer of outstretched GBP molecules the micellar state to the membrane. In reconstitution experiments with bacterial LPS, Nb77 does not visibly disrupt preformed micellar assemblies of farnesylated GBP1, yet strongly inhibits their conversion into LPS-associated coats. Productive coat formation therefore appears to require passing through an Nb77-sensitive state that is not accessible within the stable micellar assembly. This could involve local disassembly or release of smaller GBP1 species before membrane association. Our data do not distinguish between these possibilities, nor establish that micellar assemblies are obligatory precursors of coat formation.

Once GBP1 coats have formed, they become resistant to Nb77-mediated disruption. This may reflect stabilisation of the dimeric state by higher-order lateral interactions or increased avidity upon membrane association, or both. Previous observations that GDP *·* AlF_4_ ^−^-stabilised GBP1 coats remain associated after solubilisation of the underlying membrane likewise indicate substantial stabilisation within the assembled coat ^40^. Importantly, however, the experiments here capture GBP1 in a transition-state-stabilised condition. Assembly during continuous GTP turnover is likely more dynamic, and whether the same sequence of transitions occurs under physiological nucleotide turnover remains to be determined.

Nb202 recognises a composite epitope spanning the LG-domain dimer interface and selectively binds nucleotide-stabilised GBP1 dimers. Its robust recruitment to preassembled coats therefore indicates that this interface remains formed and accessible in at least a significant population of coat-associated GBP1 molecules. This observation is consistent with structural models in which the outstretched crossover dimer constitutes the main building block of the coat^19,25^ and is less readily reconciled with a coat composed of asymmetric GBP1 dimers in which this interface is dissociated or substantially rearranged in the coat-associated state ^16^. Our observations constrain differences between structural models of GBP1 assemblies, but do not quantify the fraction of GBP1 molecules adopting this conformation. The robust recruitment of Nb202 to GBP1-positive intracellular bacteria further suggests that the recognised dimeric state is abundant and accessible in the cellular context. Conversely, Nb77 reduces GBP1 coating in infected cells, consistent with its ability to interfere with productive coat assembly *in vitro*. However, the paralog specificity of the nanobodies has not been established across all seven human GBPs or mixed GBP assemblies. Thus, Nb202-positive structures in infected cells cannot yet be assigned exclusively to GBP1 in the presence of endogenous paralogs. In addition, the intrabody experiments relied on transient expression of GFP-tagged nanobodies, resulting in variable expression levels between cells and limiting quantitative comparison of coat dynamics. Inducible, genome-integrated intrabodies should provide more controlled expression and allow quantitative analysis of GBP coat formation and maturation in live cell experiments.

The different epitopes represented within the nanobody set may provide additional ways to dissect GBP coat architecture. Nb193, Nb195, Nb196 and Nb202 all preferentially recognise GBP1 at the LG-domain dimer interface, but engage this region from different orientations in ways that are likely to affect compatibility with higher-order packing in assembled coats ^19,25^. Whereas Nb202 leaves much of the lateral surface of the dimer accessible, the binding geometries of Nb193, Nb195 and Nb196 could interfere with interactions involved in higher-order packing. Their effects on membrane-associated assemblies remain to be tested, but comparison of these binders may provide a means to perturb individual interfaces without abolishing membrane recruitment altogether. Such state-selective perturbations may also provide a route to connect GBP coat architecture to downstream immune function. GBP1 can remodel and fragment LPS-containing membranes ^19^, and recent work demonstrates that GBP1-dependent deformation of bacterial and LPS membranes generates positively curved regions that promote caspase-4 binding and activation ^20^. The nanobodies described here could therefore be used to ask whether particular stages of GBP1 assembly and organisation are required for the generation of signalling-competent LPS structures.

Beyond these mechanistic insights, the nanobodies provide general tools to manipulate and visualise GBP assemblies in cells. Inhibitory nanobodies such as Nb77 or Nb75 allow acute perturbation of GBP1 function in coat formation without genetic deletion, whereas coat-compatible nanobodies such as Nb202 can label GBP-positive structures without modifying GBP1 with fluorescent protein fusions. Extending this strategy to paralog-selective binders could help resolve the composition and maturation of heterogeneous GBP coats, identify co-recruited paralogs or accessory factors, and distinguish their recruitment at different stages of assembly. Together, these reagents provide the foundation for a broader toolkit to analyse the composition and maturation of GBP coats.

## DATA AND CODE AVAILABILITY

Cryo-EM density maps of Nb193, Nb195, Nb196 and Nb202 are available in the Electron Microscopy Data Bank under accession codes EMDB-59826, EMDB-X59827, EMDB-59828, EMDB-59825. Atomic models have been deposited in the Protein Data Bank under accession codes 000034GI and 000034GJ. Nanobody sequences are available at the NanoSaurus repository under entry IDs SD-GJPK, SD-76SO, SD-G72M, SD-IY9Y, SD-FO61, SD-HOXD, SD-VEB1 and SD-X1FH.

## ACKNOWLEDGEMENTS

This work benefited from support by Nanobody4Instruct and the NKI Protein Facility through the use of resources of Instruct-ERIC (PID 10799, 21325, 23926 and 24937), part of the European Strategy Forum on Research Infrastructures (ESRFI), and the Research Foundation Flanders (FWO). We thank Alison Lundqvist and Alex Fish for technical assistance during nanobody discovery and SPR experiments. Cryo-EM data collection benefited from access to the Netherlands Centre for Electron Nanoscopy (NeCEN) with financial support from the Dutch Roadmap Grant NEMI (NWO.GWI.184.034.014). We are grateful for technical support by Willem Noteborn and Wiel Evers for cryo-EM data collection, and by Michal Shemesh for fluorescence microscopy. We thank Clémence Taisne for setting up cellular infection assays with *S*. Typhymurium. This work was supported by the European Research Council (ERC-StG-852880 and ERC-CoG-101230715 to AJ) and the Dutch Research Council (NWO.VI.C.252.143 to AJ).

## AUTHOR CONTRIBUTIONS

L.P., T.K. and A.J. designed experiments. L.P., T.K., C.P. and M.A. cloned and purified proteins. L.P., T.K., C.P., M.A. and P.N. performed biophysical experiments. L.P., T.K. and P.N. prepared cryo-EM samples. L.P. and T.K. acquired cryo-EM data. L.P. processed and analysed cryo-EM data. L.P. and A.J. built and refined the atomic models. L.P. and C.P. performed confocal imaging, and L.P. performed cryo-EM imaging of *in vitro* GBP1 coat formation. C.P. and E.G.P. performed cellular infection assays and imaging. L.P. performed confocal image analysis. L.P. and A.J. wrote the manuscript with feedback from all authors. A.J. conceptualised and supervised the study.

## Methods

### Nanobody discovery

Single-domain antibodies (nanobodies) were generated by immunising llamas with diverse GBP1 antigens (GBP1, GBP1-farnesylated, GBP1 GDP *·* AlF_4_ ^−^, GBP1-LG and GBP1-LG GDP *·* AlF_4_ ^−^) with the goal to create nanobodies that bind diverse conformational states and epitopes of GBP1. Nanobody discovery was performed as previously described ^41^. Briefly, following immunisation, lymphocytes were isolated from blood, from which the RNA was extracted and cDNA was synthesised. Sequences encoding variable domains were amplified by PCR and cloned into the pMESy4 phage display vector to create independent phage libraries. Binders were enriched via several rounds of biopanning, followed by ELISA against the diverse GBP1 antigens to screen for individual clones. In total, we identified 114 positive clones (GBP1: 17, GBP1-farnesylated: 13, GBP1 GDP *·* AlF_4_ ^−^ : 46, GBP1-LG: 12 and GBP1-LG GDP*·* AlF_4_ ^−^ : 26) which were sequenced and classified based on their CDR3 conservation using a similarity threshold of at least 80%. CDRs were numbered according to the IGMT definition ^42^.

### Construct generation and mutagenesis

All plasmids were amplified in DH5α competent cells (Invitrogen) and isolated by plasmid preparation (PureYield Plasmid Miniprep, Promega). All constructs were verified by DNA sequencing (MacroGen). Associated vectors, constructs and primers are available in tables S4/S5.

#### Cloning of GBPs and variants

GBP1 (P32455), GBP2 (P32456) and GBP3 (Q9H0R5) were codon-optimised for *E. coli* expression and acquired in pUC57 subcloning vectors (AJLD0001, AJLD0002, AJLD0003). Following PCR amplifications using primers AJLO0002 and AJLD0004 for GBP1, AJLO0006 and AJLO0007 for GBP2 and AJLO0010 and AJLO0011 for GBP3, the GBP paralogs were subsequently cloned into vector pETM14 (AJLV0009) encoding an N-terminal His_6_-tag and HRV-3C cleavage site, using BsaI/NcoI restriction-ligation (AJLD0030: GBP1; AJLD0036: GBP2; AJLD0037: GBP3). The GBP1-LG (AJLD0217) and GBP3ΔGED (AJLD0313) constructs were prepared by PCR amplification with primers AJLO0030 and AJLO0257 for GBP1-LG and AJLO0395 and AJLO0396 for GBP3ΔGED, followed by restriction-ligation with NotI/NcoI into pETM14.

The carboxyl-terminal avi-tag for GBP1 was introduced via PCR amplification using primers AJLO0002 and AJLO0058 on the GBP1 (AJLD0001) template, resulting in pUC57 GBP1-avi (AJLD0039). The construct was cleaved with NotI/BsaI and ligated into a NcoI/NotI linearised pETM14 vector (AJLV0009) resulting in GBP1-avi (AJLD0040). For GBP2, the avi-tag was introduced via Gibson assembly. The avi-tag was amplified using primers AJLO0110 and AJLO0111. The pETM14 (ALJV009) backbone was amplified using primers AJLO0112 and AJLO0113. The construct was assembled (Gibson Assembly Kit, NEB) yielding GBP2-avi (AJLD0078).

#### Cloning of FNTA/FNTB

The farnesyl transferase subunits (FNTA/FNTB) were cloned into a pCDFDuet (AJLV0038) vector enabling coexpression with GBP variants in pETM14 vectors. FNTA (AJLD0007) was amplified with primers AJLO0023 and AJLO0024. The fragment was digested with BsmBI and cloned into multiple cloning site (MCS) 1 of pCDFDuet by NcoI/NotI restriction-ligation, resulting in pCDFDuet-FNTA (AJLD0031). Similarly, FNTB (ALJD0008) was PCR amplified with primers AJLO0025 and AJLO0026. An undesired NdeI site in FNTB was removed via QuikChange mutagenesis with primers AJLO0027 and AJLO0028, yielding the new FNTB construct (AJLD0022). The FNTB insert was digested with NdI/BsmBI and cloned in the MCS2 via NdeI/XhoI-restriction-ligation of pCDFDuet-FNTA, resulting in the final construct pCDFDuet-FNTA/FNTB (AJLD0035).

#### Cloning of intrabodies

Intrabody constructs for Nb77 and Nb202 (AJLD0326, AJLD0327) were created by cloning the respective nanobody sequences into the mammalian expression vector pcDXNSMG3 (AJLV0066) via fragment exchange cloning. The nanobody sequences (AJLD0077, AJLD0202) were first cloned in the intermediate vector pINIT_kan (AJLV0081) with primers AJLO0321 and AJLO0322 and subsequently in the expression vector pcDXNSMG3 (AJLV0066). The construct encodes a Myctag, Streptavidin-binding peptide, HRV-3C cleavage site and an N-terminal EGFP. The empty vector pcDXNSM3 (ALJV0065), without EGFP, was used as dummy DNA.

#### Site-directed mutagenesis

QuickChange site-directed mutagenesis with oligos AJLO0074 and AJLO0075 was used to introduce a single-point cysteine mutant to obtain GBP1-Q577C (AJLD0056) for site-specific maleimide labelling. For the nanobodies, we mutated Ser^7^ to cysteine using primers AJLO0326 and AJLO0327 with Nb77 (AJLD0077) and Nb202 (AJLD0202) yielding Nb77-S7C (AJLD0305) and Nb202-S7C (AJLD0306) ^32^.

### Protein expression, purification and modification

#### GBPs and variants

GBP constructs GBP1, GBP2, GBP1-LG, GBP3ΔGED and GBP1-Q577C were expressed in *E. coli* BL21 (DE3) cells. Precultures were grown overnight in LB medium supplemented with corresponding antibiotics, at 37 °C while shaking. Subsequently, cultures were either grown in TB medium at 37 °C until an OD_600_ of ~ 0.7–1.0 was reached and subsequently induced with IPTG overnight, or via autoinduction in ZYP5052 media (1:50 inoculation) for 4 h at 37 °C before lowering the temperature to 18 °C to 22 °C, while shaking. Cells were harvested by centrifugation for 30 min at 4000 *·* g at 4 °C. The cell pellet was resuspended in lysis buffer (50 mM HEPES pH 7.8, 0.5 M NaCl and 0.1% Triton X-100) and lysed by freeze-thawing. DNAse I (NEB) was added to the lysate and incubated for 1.5 h while rotating at 4 °C. Cell debris was removed by centrifugation 30 min at 30 000 *·* g at 4 °C. The supernatant was either incubated with Ni-NTA resin (Thermo Scientific) or TALON resin (Clontech) for 45 min. The Ni-NTA resin was washed with 40 column volumes (CVs) buffer (50 mM HEPES pH 7.8, 0.5 M NaCl and 0.5 mM TCEP) and 40 CVs buffer B (50 mM HEPES pH 7.8, 0.5 M NaCl, 0.5 mM TCEP and 20 mM Imidazole) and eluted with elution buffer (50 mM HEPES pH 7.8, 0.5 M NaCl, 0.5 mM TCEP and 300 mM Imidazole). TALON resin was washed with 20 CVs buffer (50 mM Na_3_PO_4_, 300 mM NaCl and 10 mM Imidazole pH 7.4) and eluted with elution buffer (50 mM Na_3_PO_4_, 300 mM NaCl and 300 mM imidazole pH 7.4). Purified protein was dialysed overnight in buffer (50 mM HEPES pH 7.4, 150 mM NaCl and 0.5 mM DTT) using dialysis membrane (Spectra/Por, ROTH) at 4 °C. Samples were further purified by size-exclusion chromatography using a Superdex 200 Increase 10/300 column (Cytiva) in the same buffer and subsequently concentrated using centrifugal filters (Amicon) before being snap-frozen and stored at −80 °C. Sample purity was assessed by SDS-PAGE (Fig. S1b).

#### Biotinylation of GBP1-avi and GBP2-avi

GBP1-avi and GBP2-avi were biotinylated during bacterial expression by transforming the corresponding constructs in C43 (DE3) cells harbouring pCDFDuet-BirA (AJLD0192). Further expression and purification is as described for the other GBP constructs. Additionally, biotin was added to the cell culture during expression with a final concentration of 0.1 mM. Biotinylation yield was quantified by SDS-PAGE band shift assays with streptavidin.

#### Farnesylation of GBP1

GBP1 was farnesylated during bacterial expression by co-expressing GBP1 and pCDFDuet-FNTA/FNTB in *E. coli* BL21 (DE3) cells largely as described for GBP variants above. Following Ni-NTA purification, farnesylated GBP1 was further isolated by hydrophobic interaction chromatography prior to size-exclusion chromatography. (NH_4_)_2_SO_4_ was added to the pooled Ni-NTA elution fractions up to a final concentration of 1 M while stirring. The sample was run over a HiTrap Butyl HP column (GE Healthcare) and washed with 30 CVs high salt buffer (1.5 M (NH_4_)_2_SO_4_, 50 mM Tris-HCl pH 8.0, 5 mM MgCl_2_ and 2 mM DTT) and eluted with a linear gradient of low salt buffer (50 mM Tris-HCl pH 8, 2 mM MgCl_2_, 2 mM DTT). The farnesylated GBP1 fractions were pooled and further purified via size-exclusion chromatography as described above.

#### Nanobodies

Nanobodies were expressed in WK6 cells as described previously, with minor adaptions ^41^. Briefly, pre-cultures were grown overnight in LB medium, supplemented with ampicillin, 2% glucose and 1 mM MgCl_2_, at 37 °C while shaking. Production cultures were further grown in TB medium (1:50 inoculation) supplemented with ampicillin, 2% glucose and 2 mM MgCl_2_, at 37 °C while shaking. Once an OD_600_ of ~ 0.7–1.0 was reached, expression was induced with 1 mM IPTG and cells further incubated at 25 °C overnight while shaking. Cells were harvested by centrifugation at 4000 *·* g for 30 min at 4 °C. Nanobodies were extracted by osmotic shock. Per liter cell culture with an OD_600_ of 25, 10 mL cold TES (200 mM Tris pH 8.0, 0.5 mM EDTA, 0.5 M sucrose) was used to resuspend the cells. After 2 h of rotating at 4 °C, triple the resuspension volume of cold TES/4 (four times diluted TES) was added and incubated for an additional 1 h. The cells were centrifuged at 15 000 *·* g for 30 min at 4 °C. The supernatant was incubated with Ni-NTA resin (Thermo Scientific) for 1 h while rotating at room temperature. On a gravity column, the beads were washed with 20 CVs phosphate buffer 1 (50 mM NaH_2_PO_4_, 1 M NaCl and 10 mM imidazole, pH 7.0) and 30 CVs phosphate buffer 2 (50 mM NaH_2_PO_4_, 1 M NaCl and 10 mM imidazole, pH 6.0). Nanobody was eluted with elution buffer (50 mM NaH_2_PO_4_, 150 mM NaCl and 300 mM imidazole, pH 7.0) and dialysed in buffer (50 mM HEPES pH 7.4 and 150 mM NaCl) before being concentrated using centrifugal filters (Amicon, 3 MWCO) and snap-frozen and stored at −80 °C.

#### Site-specific labelling

Proteins were fluorescently labeled via maleimide conjugation to CF647 (GBP1-Q577C) or CF488 (Nb77-S7C and Nb202-S7C). Briefly, protein was dialysed in labelling buffer (50 mM Tris pH 7.0, 150 mM NaCl and 2 mM MgCl_2_) overnight. Dye was added in a 1:4 protein:dye molar ratio and incubated for 1 h on ice. The fluorescent protein was separated from the non-labelled fraction using a 5 mL HiTrap desalting column (Cytiva) or a Superdex 200 Increase 10/300 column (Cytiva) equilibrated in buffer (50 mM HEPES pH 7.4, 150 mM NaCl and 0.5 mM DTT).

### Biophysical characterisation

#### SEC-MALS

Size-exclusion chromatography multi-angle light scattering (SEC-MALS) was employed to assess the binding profiles of various nanobodies with GBP1, GBP2, GBP3ΔGED and GBP1-LG in their monomeric and dimeric states. Due to limited amounts of purified GBP3ΔGED, only selected nanobodies were examined for this variant. Dimerisation of GBP variants was induced by incubation of GTP transition state components (200 µM GDP, 10 mM NaF, 300 µM AlCl_3_, 2 mM MgCl_2_ and 1 mM DTT) for at least 10 min at room temperature, prior to the addition of the nanobodies. Nanobodies were added to the GBP variants in a 2:1 molar ratio and incubated for 15 min before applying ~50 µg of the mixture to a pre-equilibrated (50 mM HEPES pH 7.4, 150 mM NaCl, 0.5 mM DTT and 1% sodium azide) Superdex 200 Increase 10/300 column (Cytiva) for SEC-MALS using a high-performance liquid chromatography (HPLC) system (Agilent) equipped with a UV detector (Agilent), a multi-angle light scattering detector (DAWN 8+, Wyatt Technology) and a differential refractometer (Optilab, Wyatt Technology) with a flow rate of 0.5 mL/min at 20 °C. Chromatograms and molar masses were analysed with the ASTRA software (v7.3.1, Wyatt Technology). We utilised a differential refractive increment of 0.185 mL/g and the measured Rayleigh scattering to calculate the molar masses. Binding of the nanobodies to the GBP variants was determined by assessing the elution profile and molar masses.

#### Mass Photometry

Mass photometry measurements were performed on a Refeyn OneMP mass photometry instrument. Glass slides (High Precision cover slips, Marienfeld) were cleaned by sequential washes of isopropanol and water, followed by 30 min bath sonication. Silicon gaskets (CultureWell gaskets, Grace Bio-labs) were placed on the slides to form imaging wells. Fresh buffer (50 mM HEPES pH 7.4, 150 mM NaCl, 0.5 mM DTT) was utilised to dilute the samples to a 10 nM to 100 nM range. Nanobodies were added with a 2:1 molar ratio to the GBP variants prior to imaging and incubated for several minutes. Data was collected in the regular acquisition view and we aimed for 3.0 × 10^3^ to 6.0 × 10^3^ binding events. Data was acquired and analysed respectively through the AcquireMP and DiscoverMP software (v2.4, Refeyn).

#### Surface Plasmon Resonance

Binding kinetics were determined by Surface Plasmon Resonance (SPR). Measurements were conducted on a BIAcore T200 (GE Healthcare) with a Sensor Chip CAP (Biotin capture Kit series S, GE Healthcare). Biotinylated avi-tag GBP1 and biotinylated avi-tag GBP2 constructs were immobilised on the sensor via streptavidin-oligonucleotide coupled capturing. Dimerisation of the GBPs was induced by incubating for 10 min at room temperature with GTP transition state components (200 µM GDP, 10 mM NaF, 300 µM AlCl_3_, 2 mM MgCl_2_ and 1 mM DTT). We utilised a single-cycle kinetics approach, with an increasing analyte concentration from the dilution series 1 nM to 256 nM. Nanobody analyte was injected at 50 µL/min for 60 s, followed by a 60 s dissociation step, with a final dissociation step of 10 min for more accurate measurement of the dissociation rate (*k*_off_). The experiments were carried out in running buffer (50 mM HEPES pH 7.4, 150 mM NaCl, 0.5 mM TCEP, 0.005% (v/v) Tween-20) at 25 °C. After every run, the sensor chip was regenerated with regeneration buffer (6 M guanidine hydrochloride, 250 mM NaOH). The reference flow cell was used as a baseline. Kinetic parameters were calculated using Anabel ^43^ as single-cycle kinetics, and plotted as multi-cycle kinetics for easier interpretation. Signal drift correction was applied on the monomeric GBP2 measurements.

### Confocal microscopy of Giant Unilamellar Vesicles

#### GUV preparation

Giant unilamellar vesicles (GUVs) were prepared via PVA-assisted swelling. On a clean glass slide, 100 µL 5% PVA was evenly spread and dried at 50 °C for 45 min. A lipid mix was prepared by combining 12 µL 25 mg/mL BPLE (Avanti), 4 µL 10 mg/mL DSPE-PEG-Biotin (Avanti) and 10 µL 0.1 mg/mL Texas RED DHPE (Invitrogen) in excess chloroform. The lipid mix was spread on the PVA-coated surface and dried. Next, 250 µL of buffer (50 mM sucrose, 150 mM NaCl, 50 mM HEPES) was dispensed on-top of the dried lipid layer and incubated while gently shaking for 15 min at room temperature. Next, 30 µL GUVs were incubated with 3 µL 0.7 mg/mL farnesylated GBP1 and 2 µL 0.4 mg/mL farnesylated GBP1-CF647 together with transition state components (200 µM GDP, 10 mM NaF, 300 µM AlCl_3_, 2 mM MgCl_2_ and 1 mM DTT) for 30 min at room temperature. Imaging slides (µ-Slide 15 Well 3D, Ibidi) were briefly washed with buffer (50 mM HEPES pH 7.4, 150 mM NaCl and 0.5 mM DTT) and then sequentially washed and incubated with 1 mg/mL BSA/BSA-Biotin and with 1 mg/mL Neutravidin, each for 5 min. Lastly, 5 µL GBP1-coated GUVs was added and immobilised in each imaging well.

#### Confocal microscopy and analysis

Images were acquired on a Nikon A1R confocal microscope using a Nikon SR Apo TIRF 100*×* oil/1.49 NA objective and operated with NIS elements. The 488 nm, 561 nm, 640 nm lasers were used for excitation of the fluorophores. The fluorescent nanobodies were added directly before image acquisition. Images were analysed with FIJI and custom scripts. Briefly, a Hough Circle transform was used to identify circular GUVs in the first time point. Next, each GUVs was tracked per time frame using the lipid channel. Then, for each fluorescent channel, the maximum intensity of the radial profile was measured and divided by the mean intensity of the background surrounding the GUV, resulting in the SNR.

### Cell culture and confocal microscopy of intrabodies

#### Cell culture and infection

HeLa ΔGBP1 Tet-mCherry-GBP1 cells ^19^ were grown in RPMI-1640 medium (Gibco) with 10% FBS (Sigma) at 37 °C and 5% CO_2_ on glass cover slips. Doxycycline (Dox) was used to induce mCherry-GBP1 expression for 24 h. Cells were then further transfected with either 0.5 µg DNA of dummy DNA and GFP-Nb202, dummy DNA and GFP-Nb77 or solely dummy DNA (control) via a mixture of PEI and optiMEM (Gibco) for 4 h, before being washed with media and incubated for another 48 h.

*S*. enterica serovar Typhimurium expressing mCerulean3 was grown in TSB, overnight at 37 °C. The OD_600_ was measured and resuspended to in RPMI-1640 to an OD_600_ of 2.5. The HeLa cells were washed with PBS and 50 µL of fresh *S*. Typhimurium expressing mCerulean was added and spinoculated for 10 min at 200*·* g. Next, 45 min post infection, the cells were washed 5 times with medium. Next, 2 h post infection the cells were washed with PBS and fixated by incubating for 15 min with 4% paraformaldehyde (PFA) in PBS, before being washed again with PBS. The glass cover slides were mounted on glass slides with glycergel mounting medium (DAKO) overnight at 4 °C.

#### Confocal microscopy and analysis

Confocal imaging was performed on a LSM 900 Cryo Airyscan2 (ZEISS) in airyscan mode, using a 100*×* oil 1.4 NA objective and operated with the ZEN software. The 561 nm laser was used to image the mCherry-GBP1, the 488 nm laser for the GFP nanobodies, and the 405 nm laser for the *S*. Typhimurium expressing mCerulean. Images analysis was performed in FIJI using custom scripts. Cells showing fluorescence for both mCherry-GBP1 and GFP-Nb77/GFP-Nb202 were chosen for analysis. The coat coverage was calculated by manually tracing the circumference of individual bacteria and dividing the mCherry-GBP1 coated region by the total circumference of the bacteria. Significance was determined via a Holm-Bonferroni test.

### Cryo-EM

#### Nanobodies and GBP1 coat assembly

Small unilamellar vesicles (SUVs) were prepared by diluting 40 µL 25 mg/mL BPLE (Avanti) in 1 mL chloroform and drying the lipid in a glass vial under a nitrogen stream while rotating. The lipid was further dried with a desiccator for 1 h. Next, 1 mL of buffer (50 mM HEPES pH 7.4, 150 mM NaCl and 0.5 mM DTT) was used to resuspend the lipids, followed by vortexing and sonicating for 30 min before extruding the SUVs with a 0.05 µm cutoff. Farnesylated GBP1 was incubated with transition state components (200 µM GDP, 10 mM NaF, 300 µM AlCl_3_, 2 mM MgCl_2_ and 1 mM DTT). Next, 10 µL of a 2.4 mg/mL solution of GBP1 was either first incubated with the nanobody in a 1:2 molar ratio or with 10 µL SUVs, for 30 min. Next, the remaining SUVs and nanobody were added respectively. Cryo-EM grids were prepared by applying 3.6 µL on copper grids (QF-2/2, Cu300), blotting excess liquid for 8 s, before being plunge-frozen in liquid ethane with a Leica GP2, operated at 99% humidity and 20 °C. Per condition, 50~ micrographs were acquired on a Titan Krios (Thermo Fisher Scientific) operated at 300 kV, using a K3 Summit (Gatan) detector at a magnification of 26 000 *×* (pixel size of 3.28 Å), with an defocus ranging from −2.0 µm to −3.5 µm. Image acquisition was done via the EPU software. Movies were processed in CryoSPARC (V4.7.1) by patch-motion correction and CTF-estimation. The effect of nanobodies on GBP1 assembly on SUVs was assessed by counting the GBP1-coated and non-coated SUVs in a blinded setup. Similarly, the effect of nanobodies on GBP1-coated LPS-EB (InvivoGen) was examined. Farnesylated GBP1 was first incubated with transition state components for 15 min. Subsequently, GBP1 was incubated for another 15 min with either LPS-EB with a 10:1 weight ratio or the nanobody with a 1:2 molar ratio, before adding the remaining component. Cryo-EM grids were prepared as specified above on QF-1.2/1.3, Cu300 grids. Micrographs were acquired via the SerialEM software on a JEM-3200FSC (JEOL) operated at 300 kV, equipped with a K2 Summit (Gatan) and a in-column Ω energy filter, at a magnification of 10,000x corresponding to a pixel size of 3.7 Å.

#### Single-particle analysis

Samples (GBP1 + Nb193, GBP1 + Nb195, GBP1 + Nb196 and GBP1 + Nb202) for single-particle analysis were prepared by incubating GBP1 with transition state components (200 µM GDP, 10 mM NaF, 300 µM AlCl_3_, 2 mM MgCl_2_ and 1 mM DTT) and with a 2 to 3 molar excess of the respective nanobody for 15 min at room temperature. For the GBP1 + Nb202 sample, size-exclusion chromatography was further applied as described above, to isolate the GBP1 dimer peak with bound Nb202. Next, 3.6 µL of the respective samples were applied to glow-discharged copper grids (QF-1.2/1.3, Cu300/400) and blotted for 8 s to 10 s on a Leica GP2, operated at 99% humidity before plunge freezing in liquid ethane. Data were acquired on a JEM-3200FSC (JEOL) operated at 200 kV or 300 kV, equipped with a K2 Summit (Gatan) and a in-column Ω energy filter.

Image were acquired with SerialEM at 30 000*×* and 40 000 *×*magnification and with an underfocus range of −0.5 µm to −2.5 µm and electron fluence of ~ 60 e^-^/Å^2^. For further details see table S2. Single-particle analysis was performed using CryoSPARC (V4.7.1). Movies were processed by patch-motion correction and patch CTF estimation. Micrographs were inspected, and poor quality images were discarded. The blob picker was utilised initially to oversample particle positions, and the particle stack was subsequently cleaned by iterative 2D classification. As the middle domain (MD) and GTPase effector domain (GED) of GBP1 are highly flexible ^19^, we could only resolve the LG domains. Next, particles from 2D class averages with recognisable features were utilised to train Topaz and pick particles. Further iterative 2D classification was used to clean up particles. Abinitio reconstruction was used to obtain initial 3D classes. Subsequently, homogenous refinements, non-uniform and local refinements were used to improve the initial models. The reconstructions with Nb193, Nb195 and Nb196 could not be resolved to high resolution due to low particle numbers. As we observed anisotropy and 3D refinement artefacts occurring during processing, we advise cautious and limited interpretation of these density maps. For Nb202, we could resolve the complex structure to a global resolution of 4.1 Å after iterative 3D refinements and per-particle CTF correction. While the final map still retained a high degree of anisotropy due to preferred orientation, this could partially be mitigated via local sharpening with LocScale2^44^. Maps were processed in UCSF ChimeraX for visualisation ^45^.

#### Model building and refinement

The coordinate model for Nb202 was predicted using AlphaFold2^46^ via Colabfold ^47^. The coordinate model for the GBP1 dimer was taken from PDB:8CQB^19^. The chains were initially docked into the cryo-EM map using <monospace>phenix.emplace_local</monospace> ^48^, as implemented in UCSF ChimeraX^45^. The docked model was used as the starting model for refinement with <monospace>phenix.rocket</monospace> ^49^ using the LocScale-FEM map as the guiding potential. The resulting model was further refined with self-restraints generated at the 4.3 Å cut-off in COOT^50^ and rebuilt using gapTrick ^51^ yielding an ipTM score of 0.87. Automated refinement using Servalcat ^52^ with jelly body restraints and default parameters converged in 20 cycles with no need for interactive rebuilding.

## Supplementary Information

### Supplementary Figures

**Supplementary Fig. S1.**
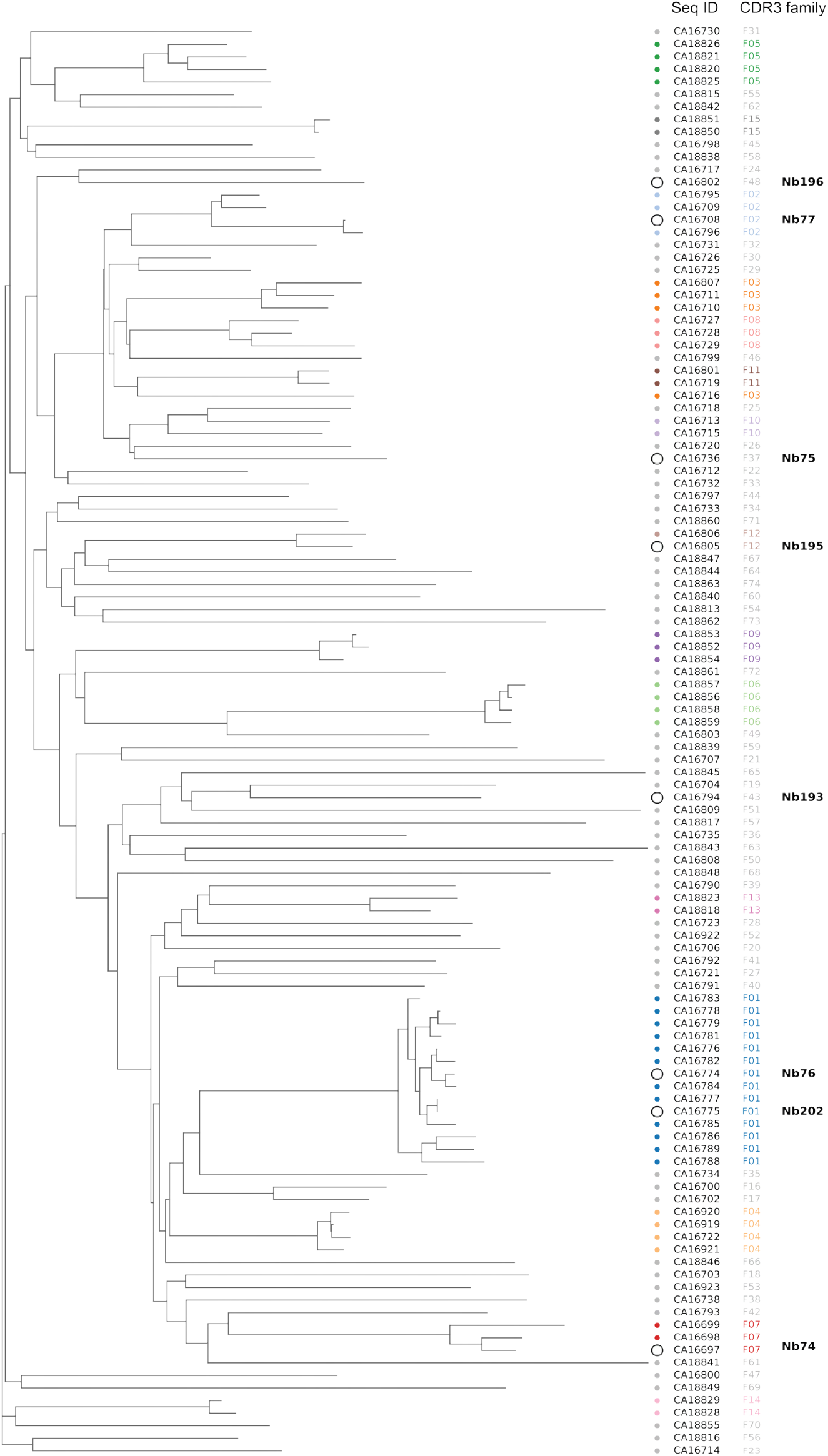
Sequence diversity and CDR3 family distribution of indentified VHHs. Neighbor-joining tree of 114 VHH sequences constructed from pairwise amino-acid p-distances calculated across the aligned VHH variable domains. Branch lengths represent amino-acid sequence divergence. The tree is shown as an unrooted sequence-similarity representation. Tips are annotated according to CDR3 sequence family. CDR3 families were defined as sequences with identical CDR3 length and >80% amino-acid identity and were grouped by single-linkage clustering, resulting in 74 families, including 59 singleton and 15 multi-member families. Multi-member CDR3 families are distinguished by colored annotations, whereas singleton families are shown separately. Nanobodies selected for detailed characterisation are highlighted and labelled with both their CA identifier and Nb designation.

**Supplementary Fig. S2.**
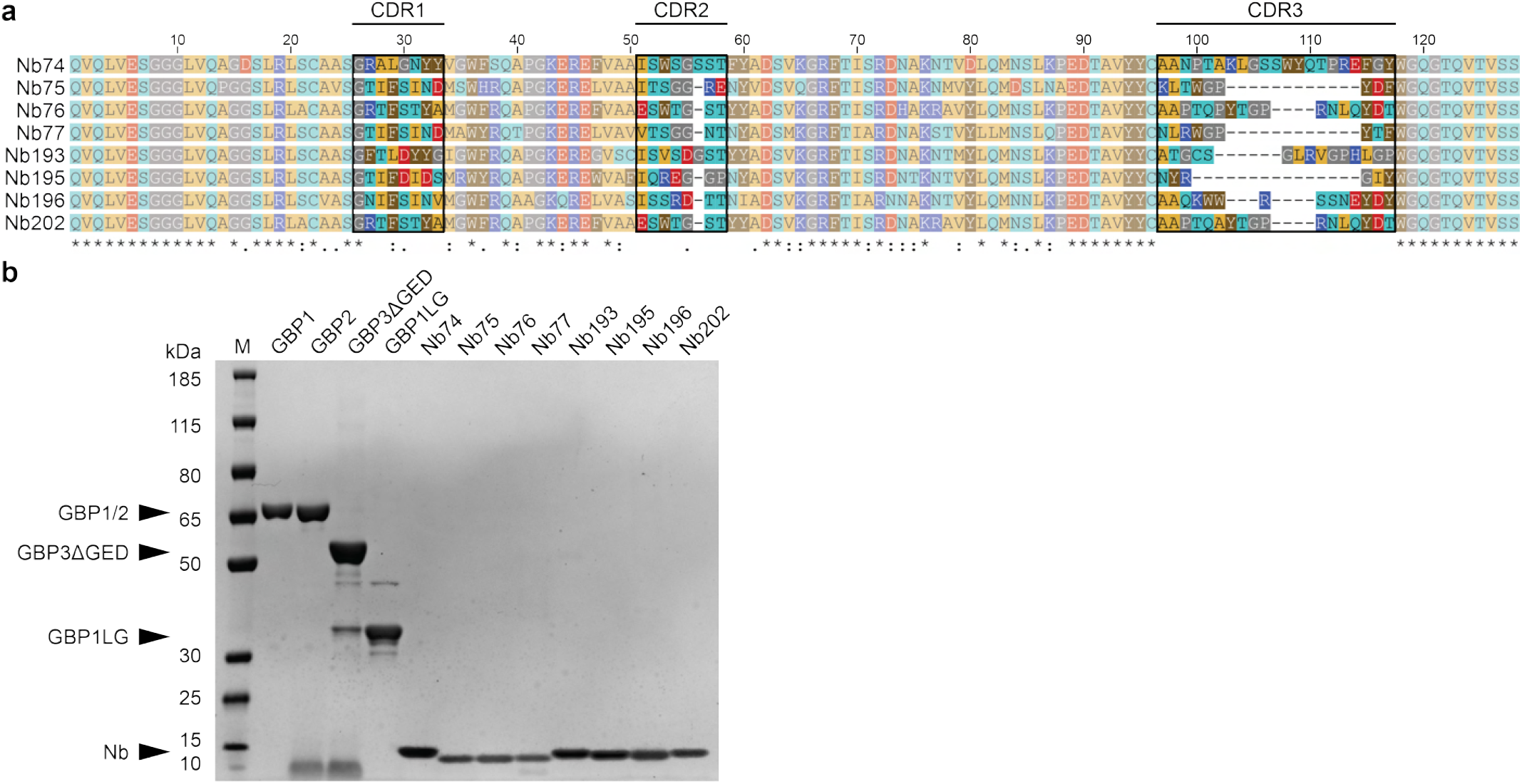
Sequence alignment of GBP-binding nanobodies. **a**, Multiple sequence alignment (MSA) of GBP-binding nanobodies. Nanobody families were classified through the conservation of the CDR3-loop (conservation >80%) according to the IMGT definition. Sequences were aligned with Clustal Omega and coloured by their biophysical properties. The symbols below the MSA follow the standard conservation consensus: (*) conserved, (:) strongly similar, (.) weakly similar and () not conserved. **b**, SDS–PAGE analysis of GBP1, GBP2, GBP3ΔGED, GBP1-LG and the selected nanobodies.

**Supplementary Fig. S3.**
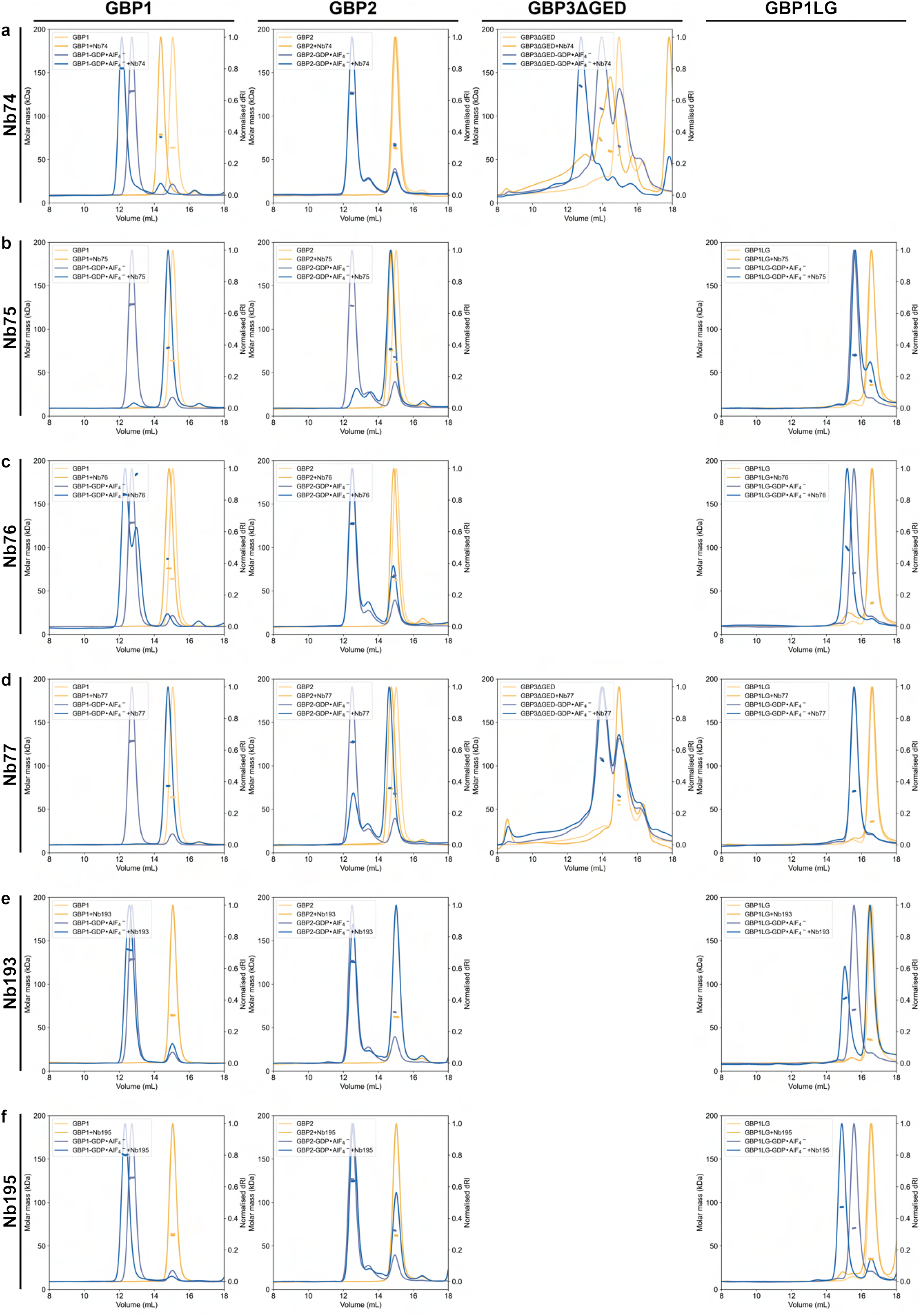
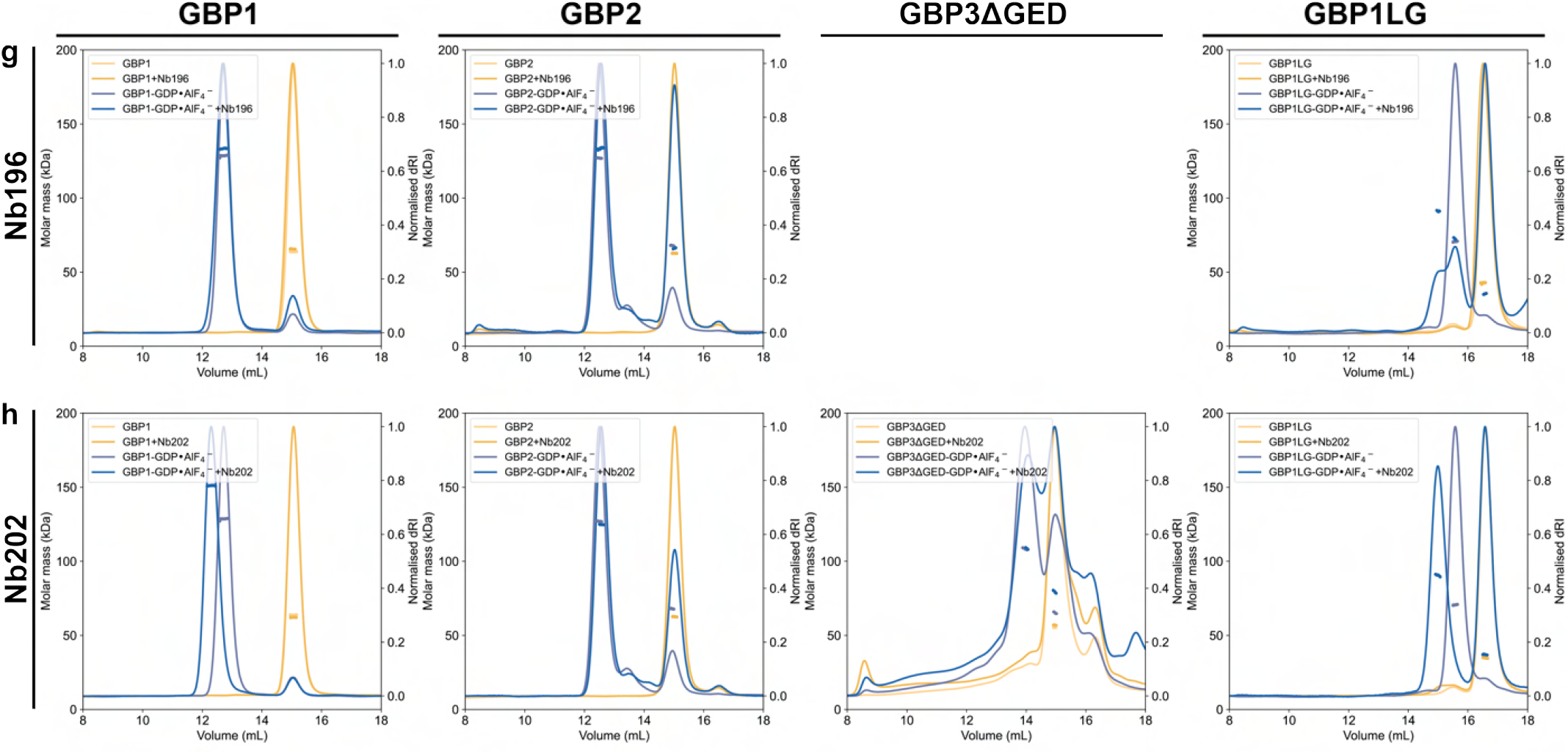
Biophysical characterisation of GBP-binding nanobodies via SEC-MALS. The binding capabilities of the nanobodies towards monomeric and dimeric GBP1, GBP2, GBP3ΔGED and GBP1-LG were examined by SEC-MALS. Light yellow and light blue represent the monomeric and dimeric GBP variants in the absence of nanobody, while the dark yellow and dark blue represent conditions with nanobody. The effect of the nanobody was assessed by comparing the calculated molar mass and shifts in elution profiles.

**Supplementary Fig. S4.**
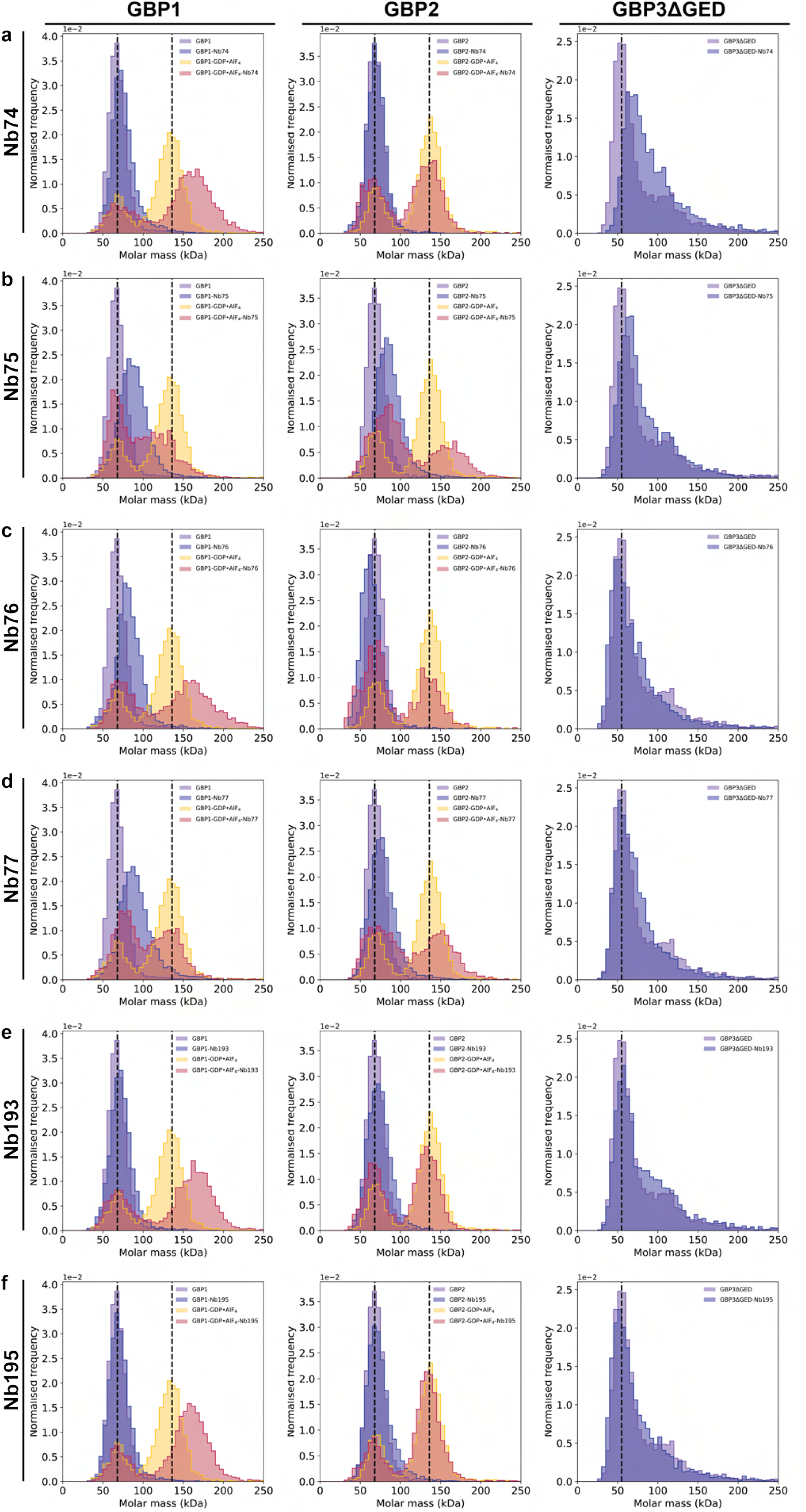
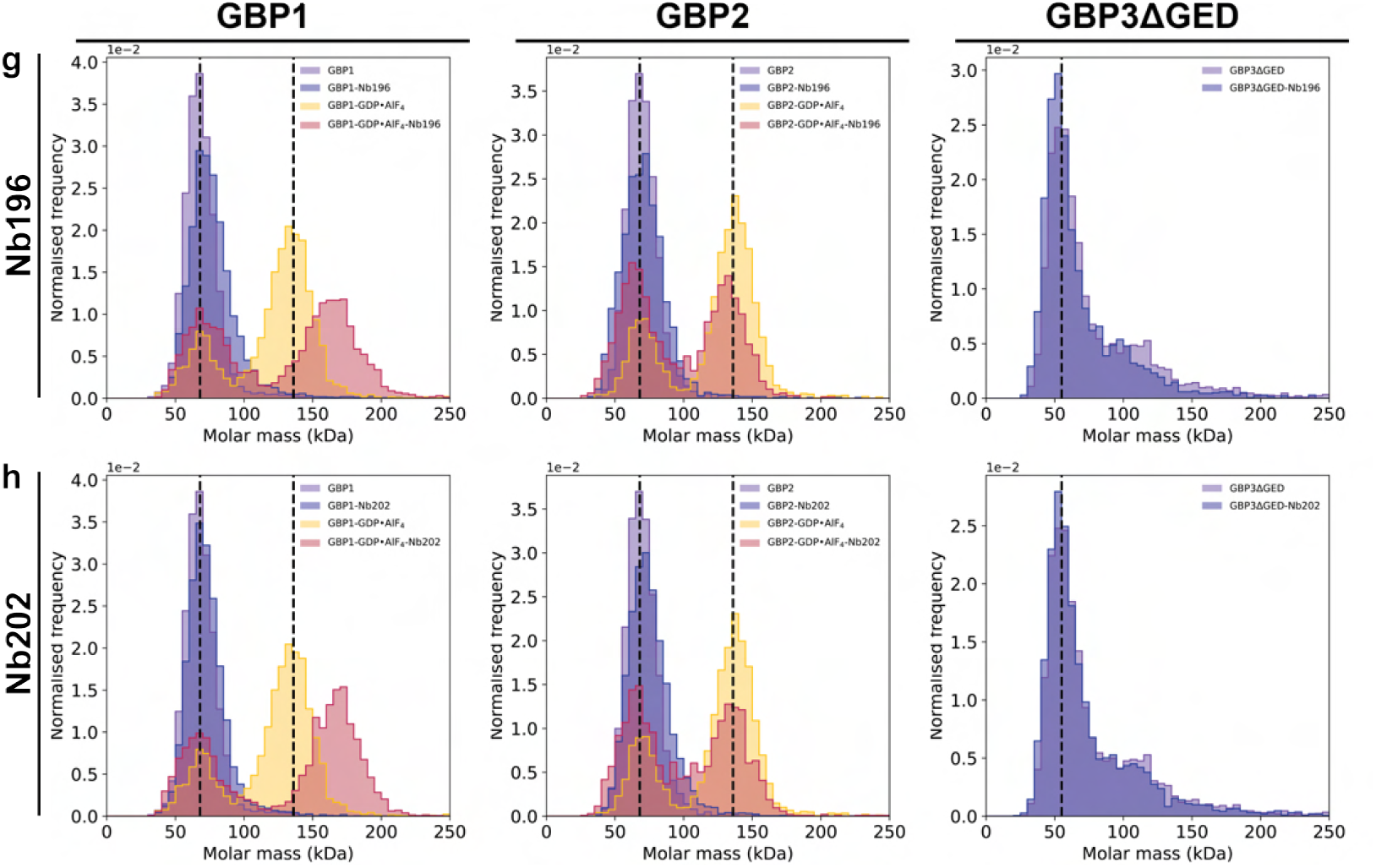
Biophysical characterisation of GBP-binding nanobodies via mass photometry. The binding capabilities of the nanobodies towards monomeric and dimeric GBP1, GBP2 and GBP3ΔGED were examined by mass photometry. Purple and yellow histograms represent the monomeric and dimeric GBP variants in the absence of nanobody, while the blue and red histograms represent conditions with nanobody. The effect of the nanobody was assessed by comparing the shift in molar mass.

**Supplementary Fig. S5.**
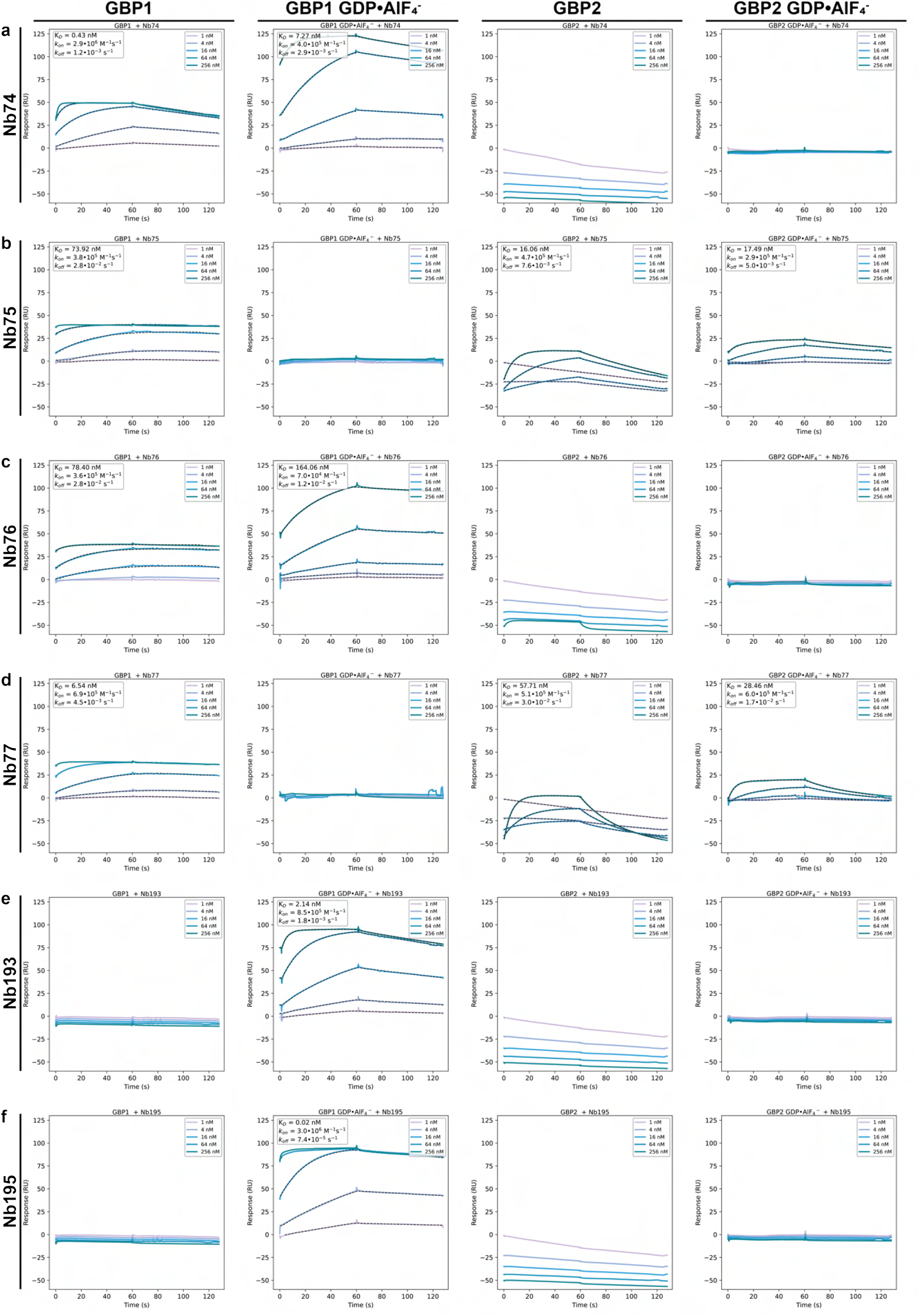
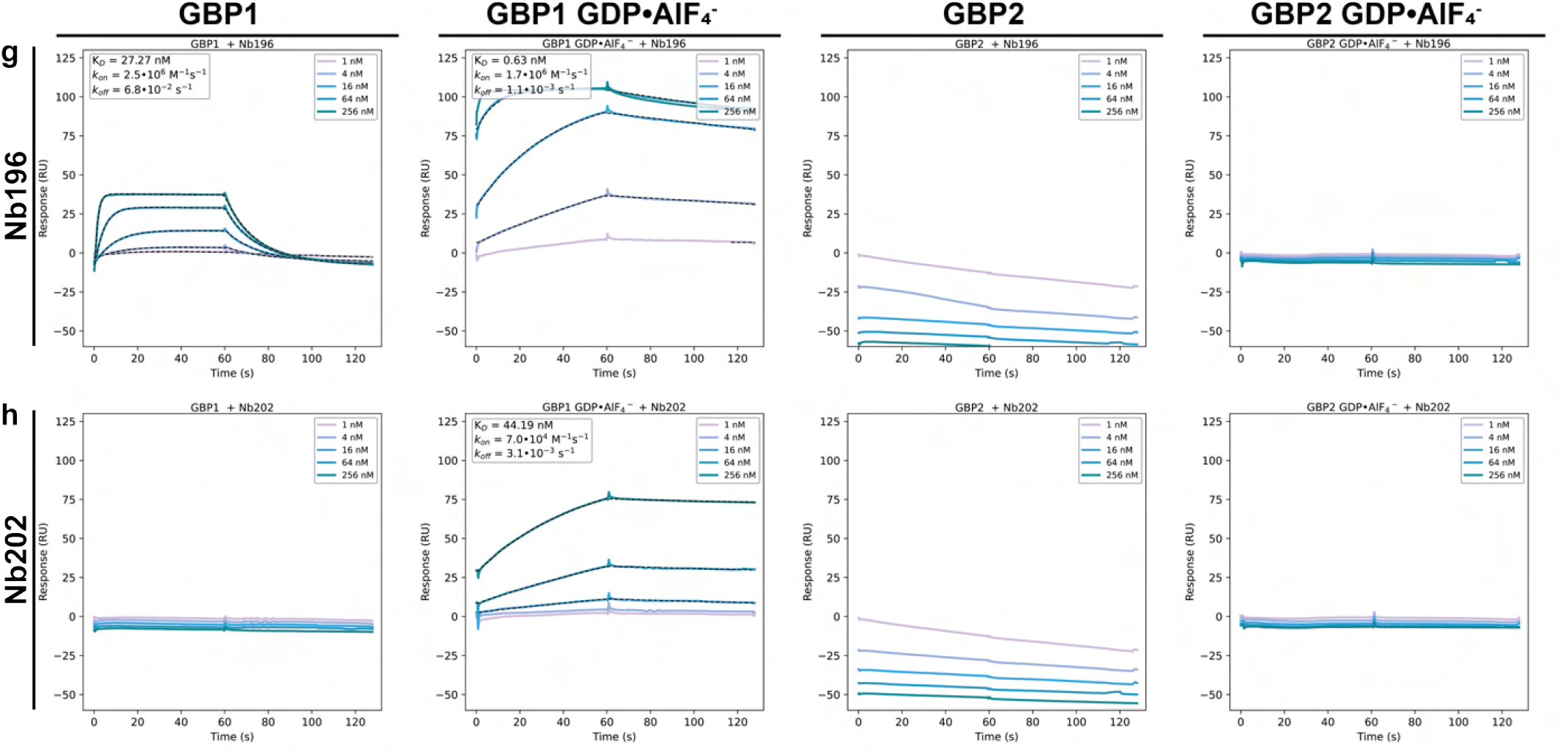
Binding kinetics of GBP-binding nanobodies measured by SPR. The binding kinetics of the nanobodies towards monomeric and dimeric GBP1 and GBP2 were measured by SPR. Single-cycle kinetics were performed with an increasing nanobody analyte (rows) concentration (inset, top right) on the immobilised GBP variant. Measurements were fitted with Anabel and calculated affinities are depicted (inset, top left).

**Supplementary Fig. S6.**
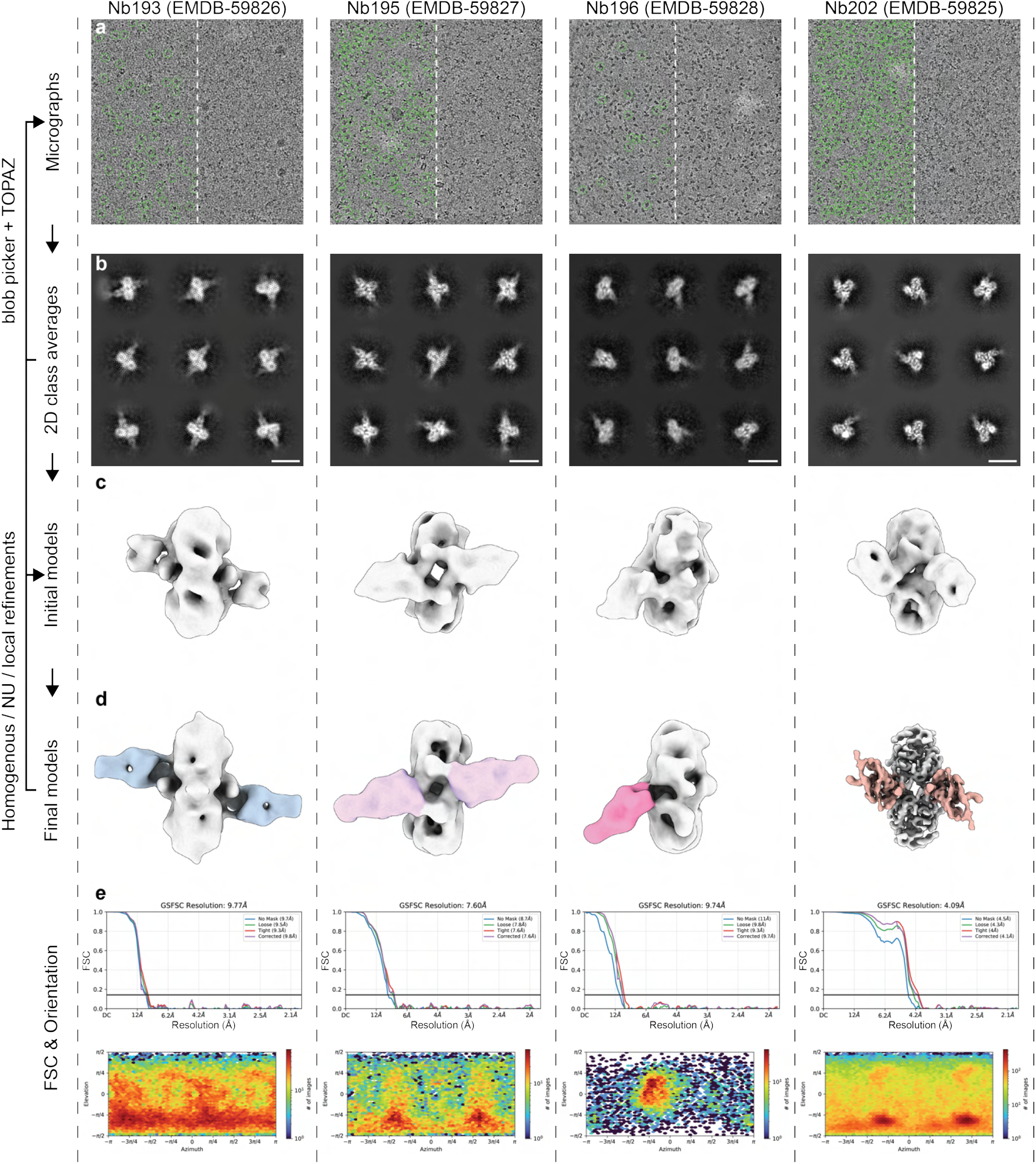
Single-particle analysis workflow. The general cryo-EM SPA workflow is depicted for Nb193 (**EMDB-59826**), Nb195 (**EMDB-59827**), Nb196 (**EMDB-598268**) and Nb202 (**EMDB-59825**) in complex with GDP *·* AlF_4_^−^ stabilised GBP1 dimers. **a**, Raw micrographs, with the final particles annotated by green circles. **b**, Selected 2D classes averages of the particles. Scale bar is 10 nm. **c**, Initial models were created by ab-initio reconstruction from the final particle sets. **d**, Iterative refinements via homogeneous refinement, non-uniform (NU) refinement and local refinement resulted in the final cryo-EM maps. **e**, The Fourier Shell correlation (FSC) and orientation maps are shown. Numerical data is provided in table S2.

**Supplementary Fig. S7.**
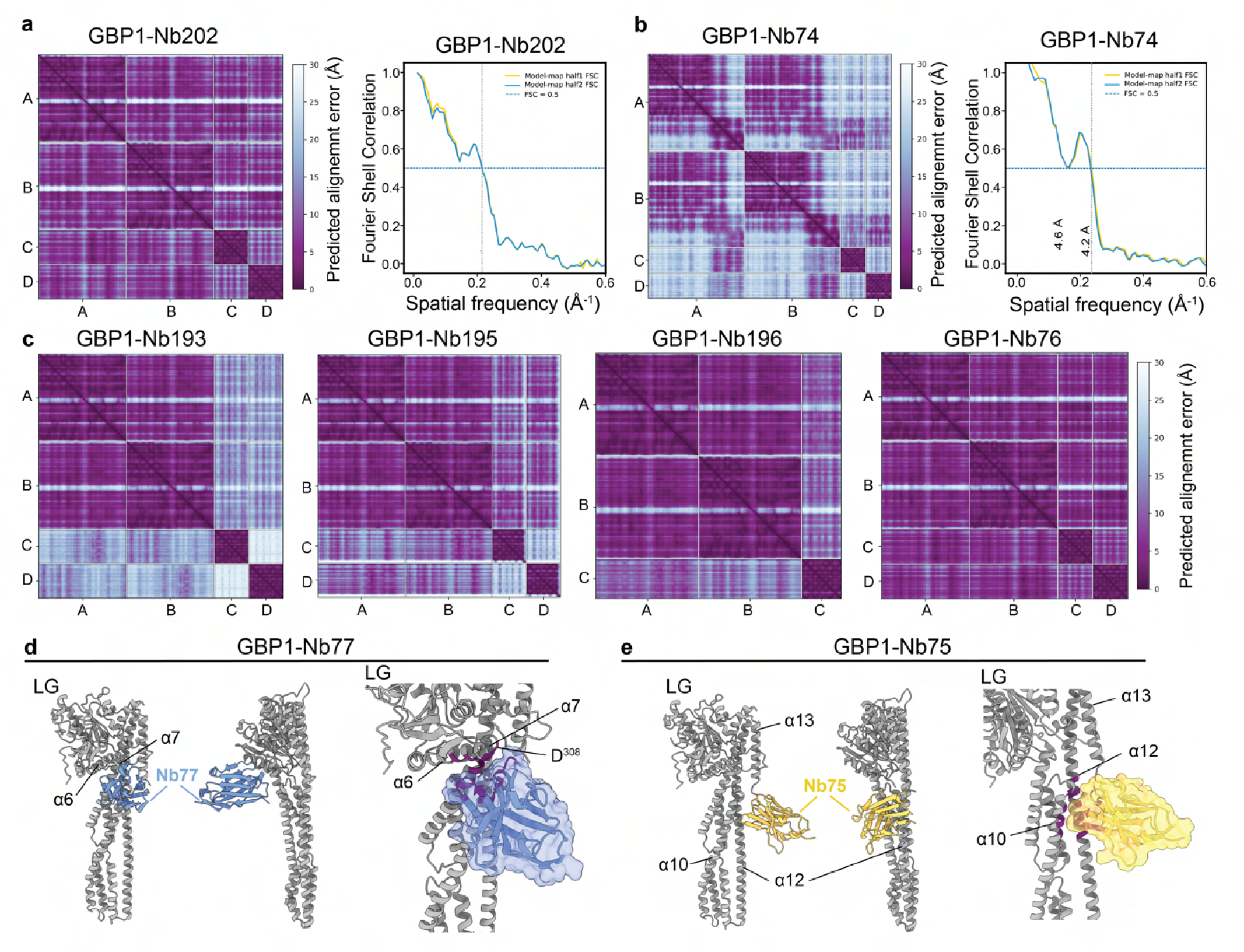
Model validation of GBP1-nanobody complexes. **a,b**, Predicted alignment error (PAE) plots of GBP1-Nb202 and GBP1-Nb74 complexes after rescoring of refined coordinate models with gapTrick using the Servalcat-refined models as template (left). Model-vs-map Fourier shell correlation curves for GBP1-Nb202 and GBP1-Nb74 coordinate models (right). **c**, PAE plots of GBP1-Nb193, GBP1-Nb195, GBP1-Nb196 and GBP1-Nb76 complexes after rescoring with gapTrick using the rigid-body docked models as template. **d,e**, Top-ranking AF3 models of GBP1-Nb75 and GBP1-Nb77. iPTM scores >0.9 indicate highly confident predictions. Nb75 is predicted to bind monomeric GBP1 at the interface formed by α10 (MD) and α12 (GED; ipTM=0.95, pTM=0.94). Nb77 is predicted to bind monomeric GBP1 at the α6-α7 loop that forms the cross-over loop in nucleotide-activated dimeric GBP1 (ipTM=0.93, pTM=0.92). In the respective close-ups residues forming contacts are highlighted in purple; the nanobody is shown in semi-transparent surface representation in these panels. We note that these models are structure predictions and need experimental validation.

**Supplementary Fig. S8.**
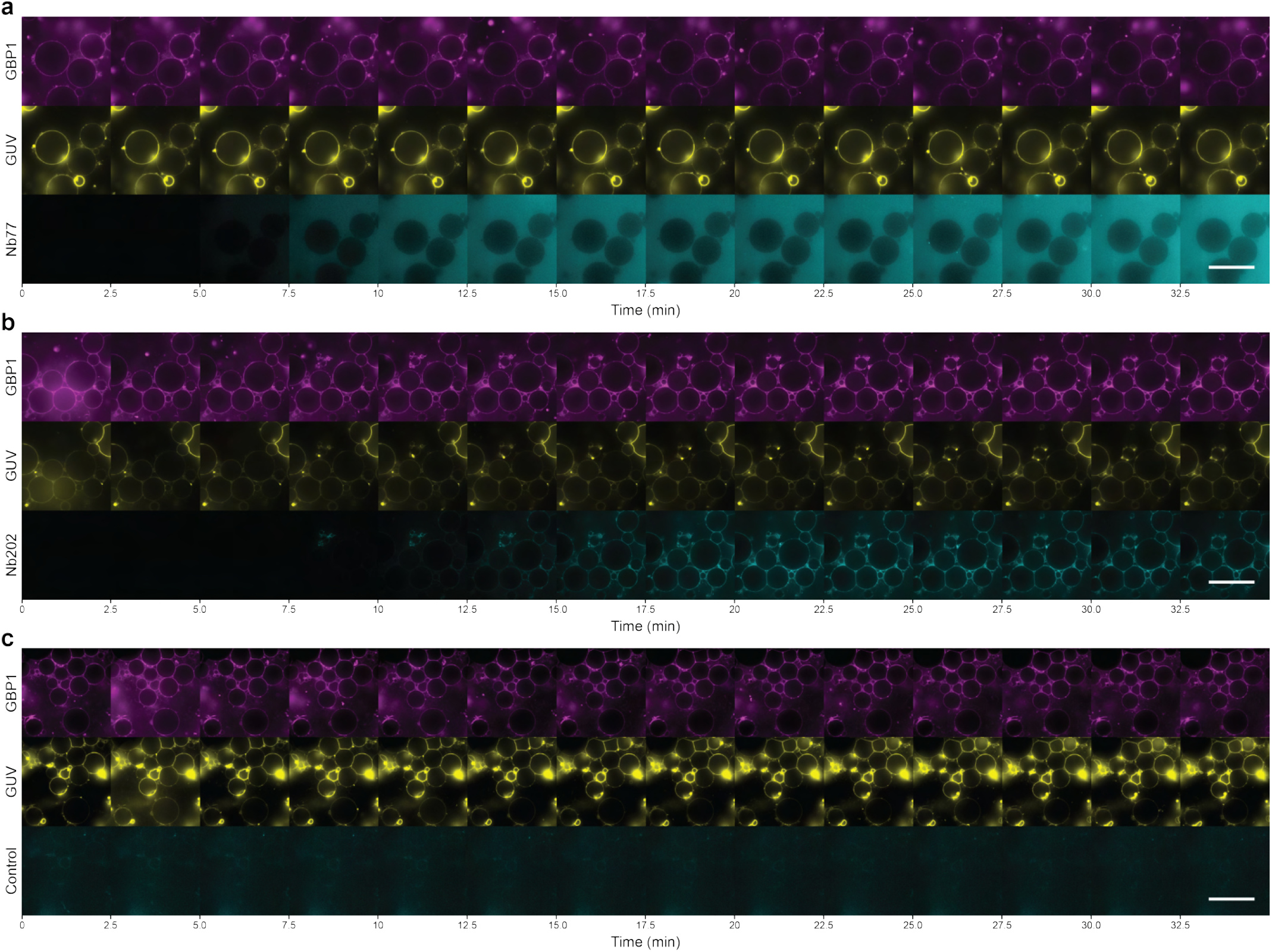
Time series of confocal imaging of nanobody recruitment to GBP1-coated GUVs. **a-c**, Representative confocal time series monitoring the association of Nb77 and Nb202 on GUVs with pre-formed GDP *·* AlF_4_ ^−^-stablised GBP1 coat. Buffer was used as a control. Scale bar is 50 µm.

**Supplementary Fig. S9.**
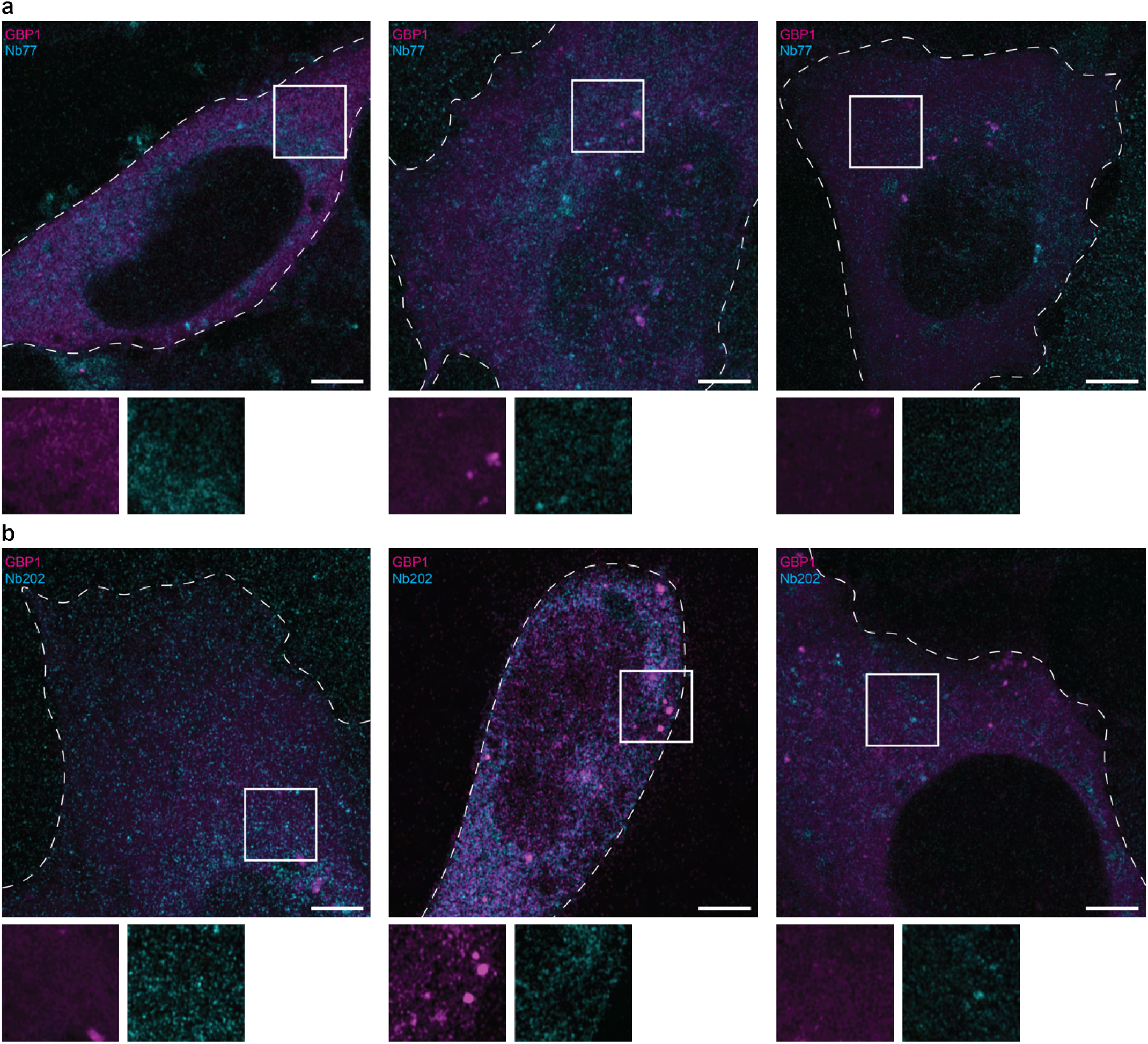
Confocal imaging of non-infected cells expressing GFP-Nb77 and GFP-Nb202. **a,b**, Confocal images of doxycycline-induced HeLa ΔGBP1 + Tet-mCherry-GBP1 cells transiently expressing GFP-Nb77 and GFP-Nb202. Lower panels show the individual fluorescence channels for the boxed regions. Scale bar is 5 µm.

**Supplementary Fig. S10.**
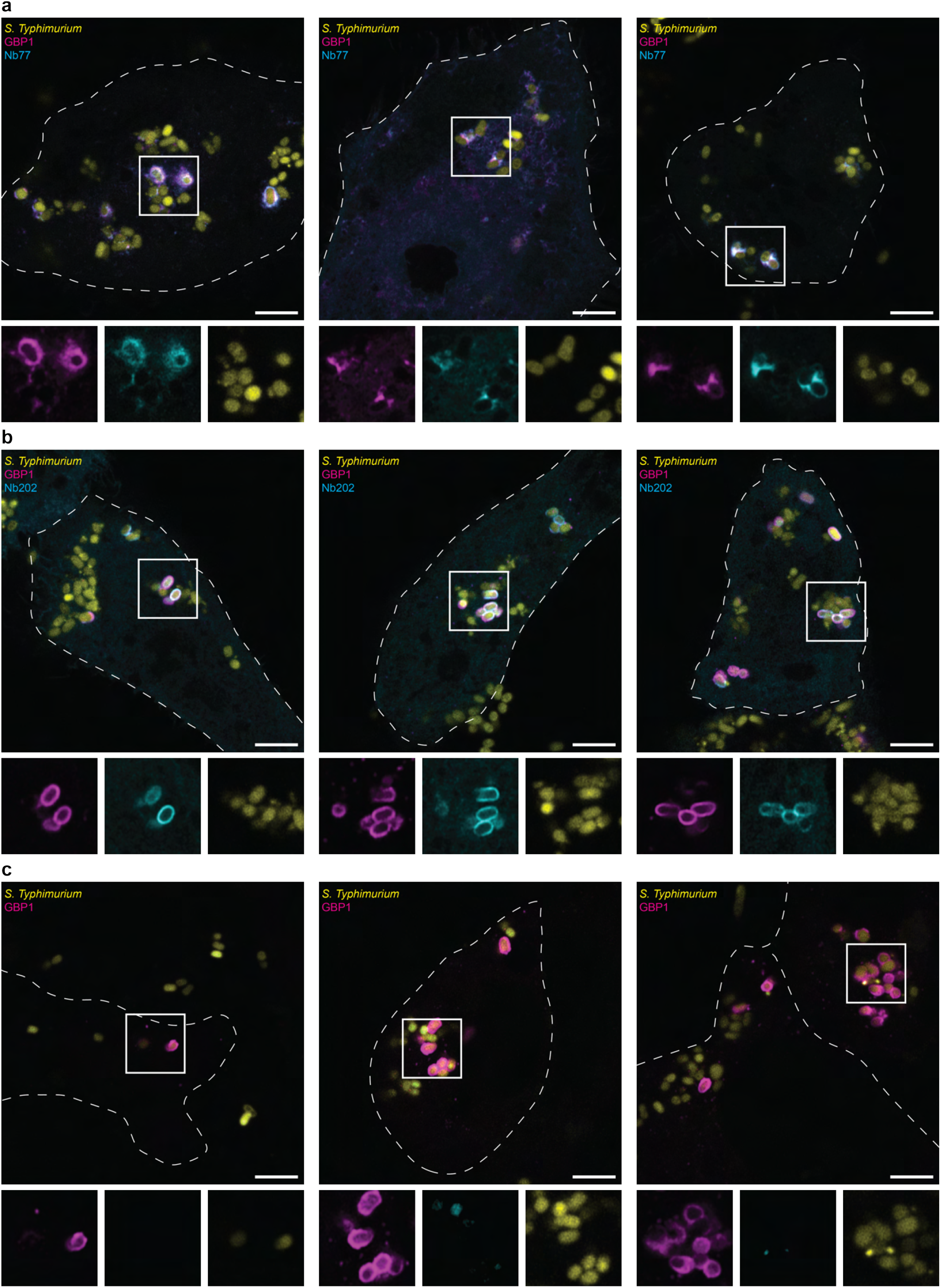
Additional examples of intrabody localisation in *S*. Typhimurium-infected cells. **a-c**, GFP-Nb77, GFP-Nb202 and dummy DNA (negative control) were transfected in HeLa ΔGBP1 + Tet-mCherry-GBP1 cells prior to doxycycline induction. Cells were subsequently infected with mCerulean3-expressing *S*. Typhimurium and imaged by confocal microscopy. Lower panels show the individual fluorescence channels for the boxed regions. Scale bar is 5 µm.

## Supplementary Tables

**Supplementary Table S1.** Kinetic parameters of GBP-binding nanobodies.

|  | GBP1 |  |  |  |  |  | GBP2 |  |  |  |  |  |
| --- | --- | --- | --- | --- | --- | --- | --- | --- | --- | --- | --- | --- |
|  | *monomer* |  |  | *dimer* |  |  | *monomer* |  |  | *dimer* |  |  |
| | $K_D$<br>(nM) | $k_{on}$<br>($M^{-1}s^{-1}$) | $k_{off}$<br>($s^{-1}$) | $K_D$<br>(nM) | $k_{on}$<br>($M^{-1}s^{-1}$) | $k_{off}$<br>($s^{-1}$) | $K_D$<br>(nM) | $k_{on}$<br>($M^{-1}s^{-1}$) | $k_{off}$<br>($s^{-1}$) | $K_D$<br>(nM) | $k_{on}$<br>($M^{-1}s^{-1}$) | $k_{off}$<br>($s^{-1}$) |
| Nb74 | 0.4 | 2.9e6 | 1.2e-3 | 7.3 | 4.0e5 | 2.9e-3 | - | - | - | - | - | - |
| Nb75 | 74.0 | 3.8e5 | 2.8e-2 | - | - | - | 16.1 | 4.7e5 | 0.0076 | 17.5 | 2.9e5 | 5.0e-3 |
| Nb76 | 78.4 | 3.6e5 | 2.8e-2 | 164.1 | 7.1e4 | 1.2e-2 | - | - | - | - | - | - |
| Nb77 | 6.5 | 6.9e5 | 4.5e-3 | - | - | - | 57.7 | 5.1e5 | 0.0296 | 28.5 | 6.0e5 | 1.7e-2 |
| Nb193 | - | - | - | 2.1 | 8.5e5 | 1.8e-3 | - | - | - | - | - | - |
| Nb195 | - | - | - | 0.02 | 3.0e6 | 1.0e-4 | - | - | - | - | - | - |
| Nb196 | 27.3 | 2.5e6 | 6.8e-2 | 0.6 | 1.7e6 | 1.1e-3 | - | - | - | - | - | - |
| Nb202 | - | - | - | 44.2 | 7.0e4 | 3.1e-3 | - | - | - | - | - | - |

**Supplementary Table S2.** Cryo-EM data collection and single-particle image analysis.

|  | **Nb193**<br>**EMDB-59826** | **Nb195**<br>**EMDB-59827** | **Nb196**<br>**EMDB-59828** | **Nb202**<br>**EMDB-59825** |
| --- | --- | --- | --- | --- |
| Specimen | GBP1 + Nb193 | GBP1 + Nb195 | GBP1 + Nb196 | GBP1 + Nb202 |
| Microscope | JEOL 3200-FSC | JEOL 3200-FSC | JEOL 3200-FSC | JEOL 3200-FSC |
| Voltage (kV) | 300 | 200 | 200 | 300 |
| Detector | K2-XP | K2-XP | K2-XP | K2-XP |
| Energy filter | In-column $\Omega$ filter | In-column $\Omega$ filter | In-column $\Omega$ filter | In-column $\Omega$ filter |
| Movies | 1191 | 195 | 436 | 597 |
| Pixel size (Å) | 0.98 | 0.94 | 0.94 | 0.98 |
| Electron exposure (e<sup>-</sup>/Å<sup>2</sup>) | 56.2 | 61.0 | 60.0 | 60.1 |
| Frames | 35 | 53 | 45 | 38 |
| Defocus range (μm) | -1.25 to -2.75 | -1 to -2.5 | -1 to -2.5 | -0.5 to -4 |
| Symmetry | C2 | C2 | C1 | C2 |
| Guinier B-factor (Å<sup>2</sup>) | N.A. | N.A. | N.A. | 196 |
| Particles | 56,885 | 21,180 | 6,449 | 143,986 |
| Nominal resolution (Å) | 9.1 | 7.5 | 9.2 | 4.1 |

**Supplementary Table S3.** Model refinement statistics.

|  | **GBP1-Nb202**<br>**000034GI** | **GBP1-Nb74**<br>**000034GJ** |
| --- | --- | --- |
| Initial model used<br>(PDB code) | **8CQB** | **8CQB** |
| Model resolution<br>(Å, FSC = 0.5) | 4.6 | 4.0 |
| CC mask | 0.764 | 0.751 |
| **Model composition** |  |  |
| Nonhydrogen atoms | 6804 | 9646 |
| Protein residues | 868 | 1222 |
| **B factors** |  |  |
| Overall (Å) | 139.2 | 75.3 |
| GBP1 (Å) | 141.9 | 72.4 |
| Nb (Å) | 138.2 | 89.2 |
| **RMSD** |  |  |
| Bond lengths (Å) | 0.013 | 0.003 |
| Bond angles (°) | 0.721 | 0.604 |
| **Validation** |  |  |
| MolProbity score | 0.92 |  |
| Clashscore | 1.85 | 3.45 |
| Rotamer outliers (%) | 0.27 | 0.75 |
| **Ramachandran plot** |  |  |
| Favored (%) | 97.8 | 96.9 |
| Allowed (%) | 2.2 | 3.1 |
| Disallowed (%) | 0 | 0 |

**Supplementary Table S4.**
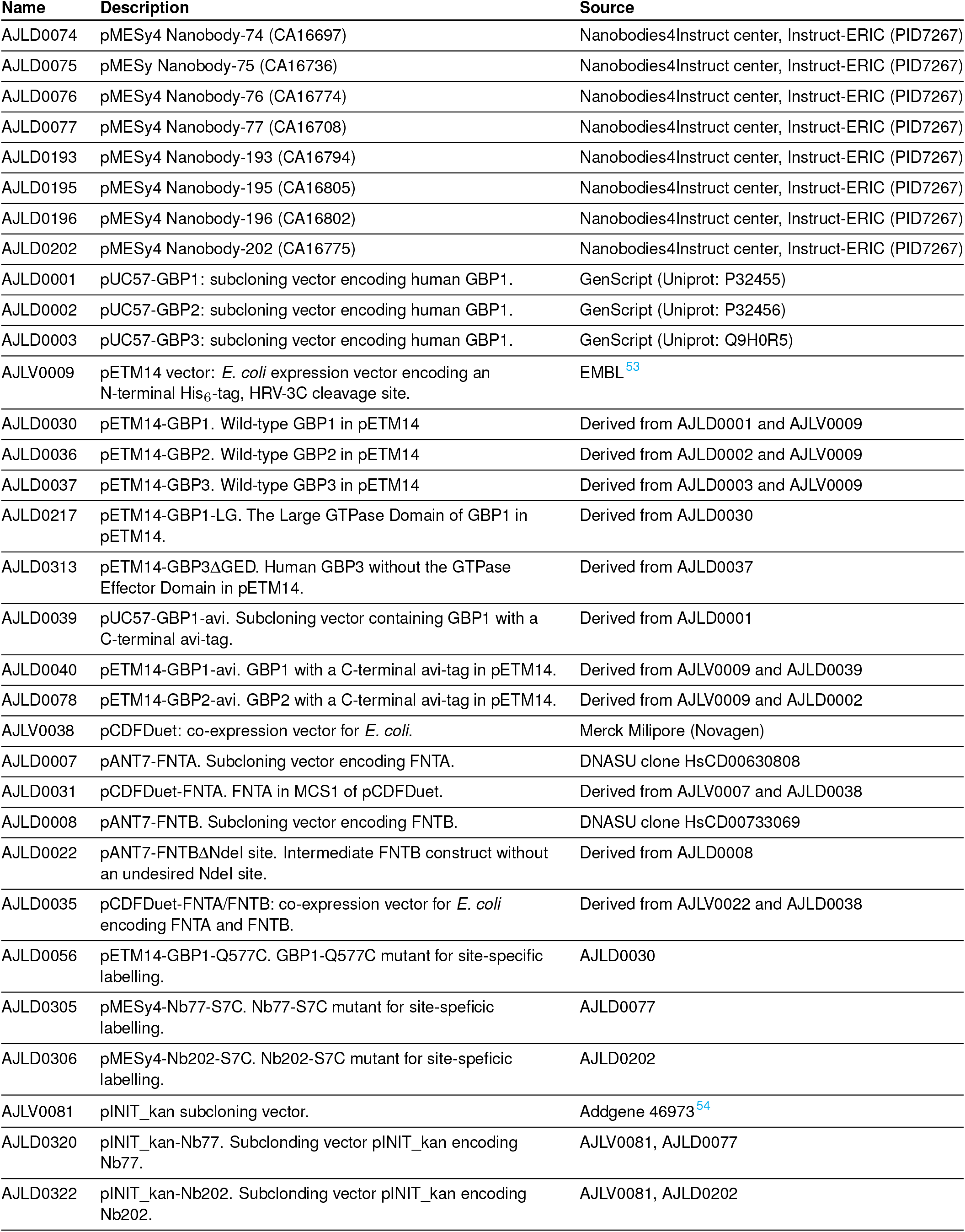
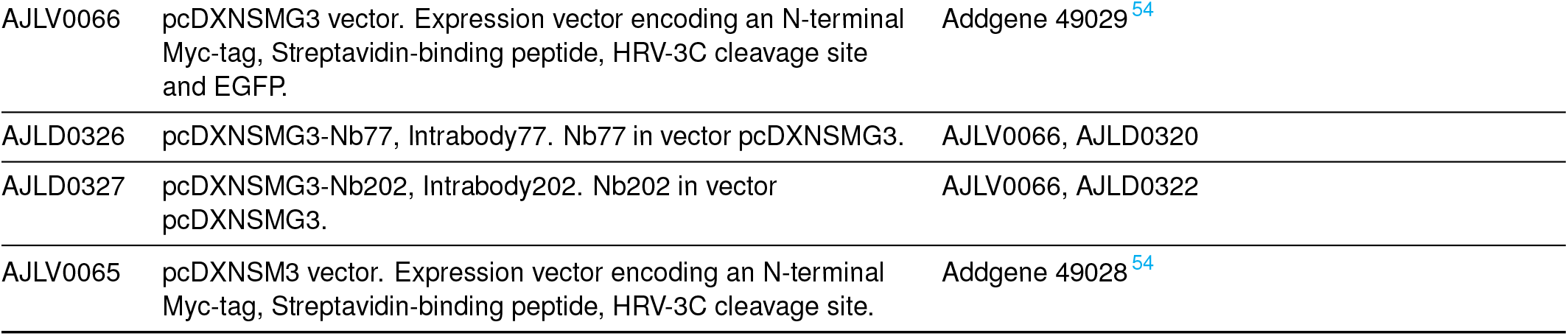
List of vectors and constructs.

| Name | Description | Source |
| --- | --- | --- |
| AJLD0074 | pMESy4 Nanobody-74 (CA16697) | Nanobodies4Instruct center, Instruct-ERIC (PID7267) |
| AJLD0075 | pMESy Nanobody-75 (CA16736) | Nanobodies4Instruct center, Instruct-ERIC (PID7267) |
| AJLD0076 | pMESy4 Nanobody-76 (CA16774) | Nanobodies4Instruct center, Instruct-ERIC (PID7267) |
| AJLD0077 | pMESy4 Nanobody-77 (CA16708) | Nanobodies4Instruct center, Instruct-ERIC (PID7267) |
| AJLD0193 | pMESy4 Nanobody-193 (CA16794) | Nanobodies4Instruct center, Instruct-ERIC (PID7267) |
| AJLD0195 | pMESy4 Nanobody-195 (CA16805) | Nanobodies4Instruct center, Instruct-ERIC (PID7267) |
| AJLD0196 | pMESy4 Nanobody-196 (CA16802) | Nanobodies4Instruct center, Instruct-ERIC (PID7267) |
| AJLD0202 | pMESy4 Nanobody-202 (CA16775) | Nanobodies4Instruct center, Instruct-ERIC (PID7267) |
| AJLD0001 | pUC57-GBP1: subcloning vector encoding human GBP1. | GenScript (Uniprot: P32455) |
| AJLD0002 | pUC57-GBP2: subcloning vector encoding human GBP1. | GenScript (Uniprot: P32456) |
| AJLD0003 | pUC57-GBP3: subcloning vector encoding human GBP1. | GenScript (Uniprot: Q9H0R5) |
| AJLV0009 | pETM14 vector: *E. coli* expression vector encoding an N-terminal His<sub>6</sub>-tag, HRV-3C cleavage site. | EMBL <sup>53</sup> |
| AJLD0030 | pETM14-GBP1. Wild-type GBP1 in pETM14 | Derived from AJLD0001 and AJLV0009 |
| AJLD0036 | pETM14-GBP2. Wild-type GBP2 in pETM14 | Derived from AJLD0002 and AJLV0009 |
| AJLD0037 | pETM14-GBP3. Wild-type GBP3 in pETM14 | Derived from AJLD0003 and AJLV0009 |
| AJLD0217 | pETM14-GBP1-LG. The Large GTPase Domain of GBP1 in pETM14. | Derived from AJLD0030 |
| AJLD0313 | pETM14-GBP3ΔGED. Human GBP3 without the GTPase Effector Domain in pETM14. | Derived from AJLD0037 |
| AJLD0039 | pUC57-GBP1-avi. Subcloning vector containing GBP1 with a C-terminal avi-tag. | Derived from AJLD0001 |
| AJLD0040 | pETM14-GBP1-avi. GBP1 with a C-terminal avi-tag in pETM14. | Derived from AJLV0009 and AJLD0039 |
| AJLD0078 | pETM14-GBP2-avi. GBP2 with a C-terminal avi-tag in pETM14. | Derived from AJLV0009 and AJLD0002 |
| AJLV0038 | pCDFDuet: co-expression vector for *E. coli*. | Merck Milipore (Novagen) |
| AJLD0007 | pANT7-FNTA. Subcloning vector encoding FNTA. | DNASU clone HsCD00630808 |
| AJLD0031 | pCDFDuet-FNTA. FNTA in MCS1 of pCDFDuet. | Derived from AJLV0007 and AJLD0038 |
| AJLD0008 | pANT7-FNTB. Subcloning vector encoding FNTB. | DNASU clone HsCD00733069 |
| AJLD0022 | pANT7-FNTBΔNdel site. Intermediate FNTB construct without an undesired Ndel site. | Derived from AJLD0008 |
| AJLD0035 | pCDFDuet-FNTA/FNTB: co-expression vector for *E. coli* encoding FNTA and FNTB. | Derived from AJLV0022 and AJLD0038 |
| AJLD0056 | pETM14-GBP1-Q577C. GBP1-Q577C mutant for site-specific labelling. | AJLD0030 |
| AJLD0305 | pMESy4-Nb77-S7C. Nb77-S7C mutant for site-specific labelling. | AJLD0077 |
| AJLD0306 | pMESy4-Nb202-S7C. Nb202-S7C mutant for site-specific labelling. | AJLD0202 |
| AJLV0081 | pINIT_kan subcloning vector. | Addgene 46973 <sup>54</sup> |
| AJLD0320 | pINIT_kan-Nb77. Subcloning vector pINIT_kan encoding Nb77. | AJLV0081, AJLD0077 |
| AJLD0322 | pINIT_kan-Nb202. Subcloning vector pINIT_kan encoding Nb202. | AJLV0081, AJLD0202 |
| AJLV0066 | pcDXNSMG3 vector. Expression vector encoding an N-terminal Myc-tag, Streptavidin-binding peptide, HRV-3C cleavage site and EGFP. | Addgene 49029<sup>54</sup> |
| AJLD0326 | pcDXNSMG3-Nb77, Intrabody77. Nb77 in vector pcDXNSMG3. | AJLV0066, AJLD0320 |
| AJLD0327 | pcDXNSMG3-Nb202, Intrabody202. Nb202 in vector pcDXNSMG3. | AJLV0066, AJLD0322 |
| AJLV0065 | pcDXNSM3 vector. Expression vector encoding an N-terminal Myc-tag, Streptavidin-binding peptide, HRV-3C cleavage site. | Addgene 49028<sup>54</sup> |

**Supplementary Table S5.**
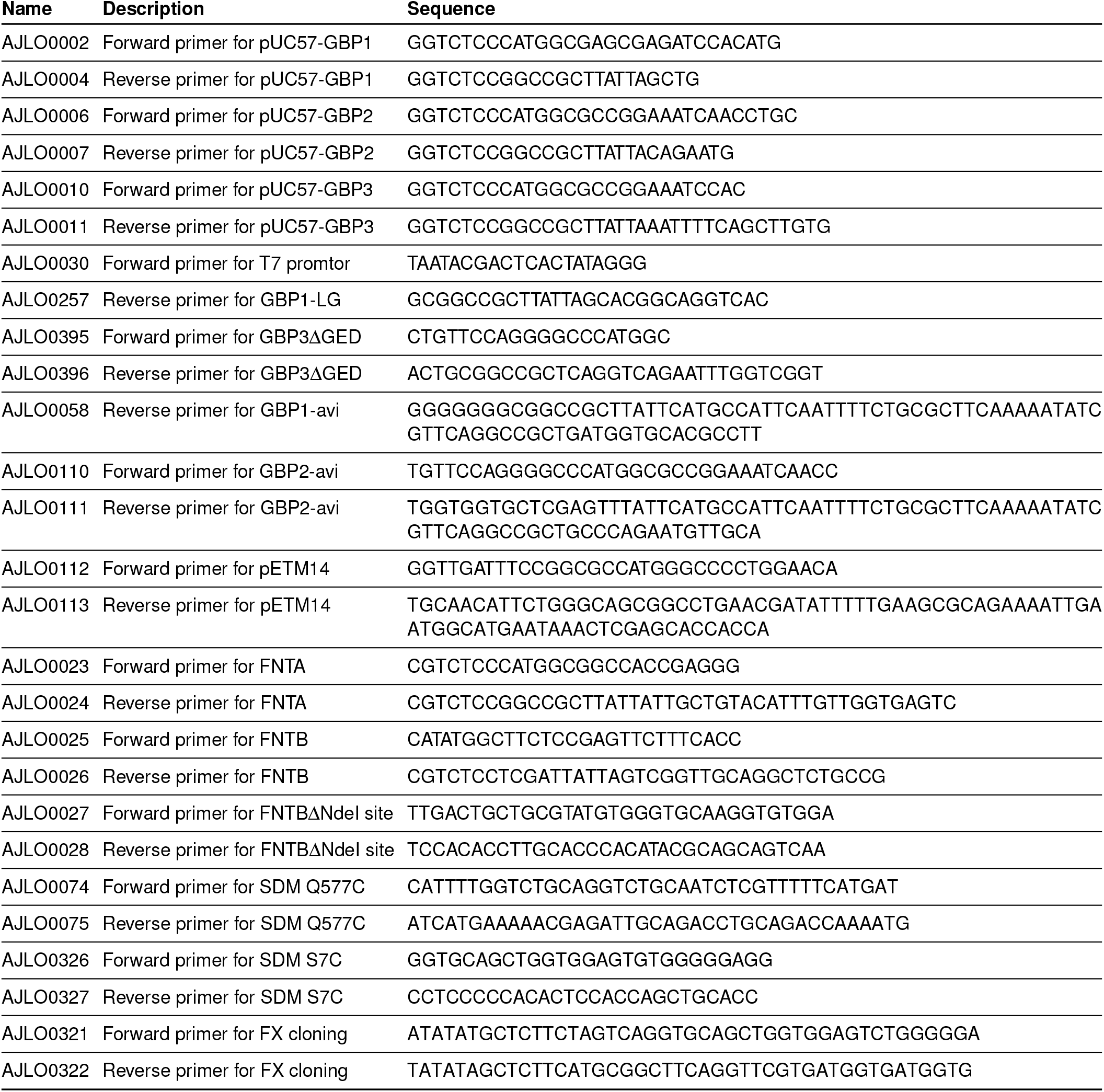
List of oligonucleotides.

**Supplementary Table S6.**
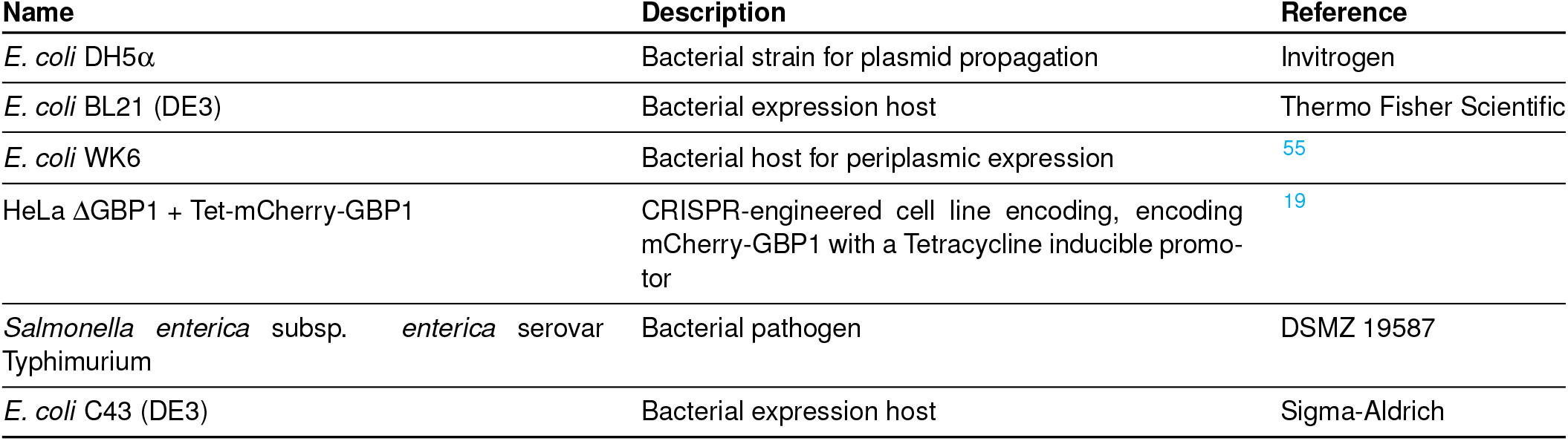
Bacterial strains and mammalian cells.

## References

1. B.-H. Kim, A. R. Shenoy, P. Kumar, R. Das, S. Tiwari and J. D. MacMicking. A Family of IFN-γ–Inducible 65-kD GTPases Protects Against Bacterial Infection. Science, 332(6030): 717–721, May 2011.

2. K. Tretina, E.-S. Park, A. Maminska and J. D. MacMicking. Interferon-induced guanylate-binding proteins: Guardians of host defense in health and disease. Journal of Experimental Medicine, 216(3):482–500, February 2019.

3. Y. Rivera-Cuevas, B. Clough and E.-M. Frickel. Human guanylate-binding proteins in intracellular pathogen detection, destruction, and host cell death induction. Current Opinion in Immunology, 84:102373, 2023.

4. M. A. Olszewski, J. Gray and D. J. Vestal. In Silico Genomic Analysis of the Human and Murine Guanylate-Binding Protein (GBP) Gene Clusters. Journal of Interferon & Cytokine Research, 26(5):328–352, May 2006.

5. S. L. Gupta, B. Y. Rubin and S. L. Holmes. Interferon action: induction of specific proteins in mouse and human cells by homologous interferons. Proceedings of the National Academy of Sciences, 76(10):4817–4821, October 1979.

6. Y. S. Cheng, R. J. Colonno and F. H. Yin. Interferon induction of fibroblast proteins with guanylate binding activity. Journal of Biological Chemistry, 258(12):7746–7750, June 1983.

7. E. Kravets, D. Degrandi, S. Weidtkamp-Peters, B. Ries, C. Konermann, S. Felekyan, J. M. Dargazanli, G. J. K. Praefcke, C. A. M. Seidel, L. Schmitt, S. H. J. Smits, et al. The GTPase activity of murine guanylate-binding protein 2 (mGBP2) controls the intracellular localization and recruitment to the parasitophorous vacuole of Toxoplasma gondii. The Journal of Biological Chemistry, 287(33):27452–27466, August 2012.

8. E. Kravets, D. Degrandi, Q. Ma, T.-O. Peulen, V. Klümpers, S. Felekyan, R. Kühnemuth, S. Weidtkamp-Peters, C. A. Seidel, et al. Guanylate binding proteins directly attack Toxoplasma gondii via supramolecular complexes. eLife, 5:e11479, January 2016.

9. D. Fisch, H. Bando, B. Clough, V. Hornung, M. Yamamoto, A. R. Shenoy, et al. Human GBP1 is a microbe-specific gatekeeper of macrophage apoptosis and pyroptosis. The EMBO Journal, 38(13):e100926, July 2019.

10. M. P. Wandel, B.-H. Kim, E.-S. Park, K. B. Boyle, K. Nayak, B. Lagrange, A. Herod, T. Henry, M. Zilbauer, J. Rohde, J. D. MacMicking, et al. Guanylate-binding proteins convert cytosolic bacteria into caspase-4 signaling platforms. Nature Immunology, 21(8):880–891, August 2020.

11. J. C. Santos, D. Boucher, L. K. Schneider, B. Demarco, M. Dilucca, K. Shkarina, R. Heilig, K. W. Chen, R. Y. H. Lim, et al. Human GBP1 binds LPS to initiate assembly of a caspase-4 activating platform on cytosolic bacteria. Nature Communications, 11(1):3276, June 2020.

12. E. Meunier, M. S. Dick, R. F. Dreier, N. Schürmann, D. K. Broz, S. Warming, M. Roose-Girma, D. Bumann, N. Kayagaki, K. Takeda, M. Yamamoto, et al. Caspase-11 activation requires lysis of pathogen-containing vacuoles by IFN-induced GTPases. Nature, 509 (7500):366–370, May 2014.

13. E. Meunier, P. Wallet, R. F. Dreier, S. Costanzo, L. Anton, S. Rühl, S. Dussurgey, M. S. Dick, A. Kistner, M. Rigard, D. Degrandi, K. Pfeffer, M. Yamamoto, T. Henry, et al. Guanylate-binding proteins promote activation of the AIM2 inflammasome during infection with Francisella novicida. Nature Immunology, 16(5):476–484, May 2015.

14. N. Britzen-Laurent, M. Bauer, V. Berton, N. Fischer, A. Syguda, S. Reipschläger, E. Naschberger, C. Herrmann, et al. Intracellular Trafficking of Guanylate-Binding Proteins Is Regulated by Heterodimerization in a Hierarchical Manner. PLOS ONE, 5(12):e14246, December 2010.

15. M. Kutsch, L. Sistemich, C. F. Lesser, M. B. Goldberg, C. Herrmann and J. Coers. Direct binding of polymeric GBP1 to LPS disrupts bacterial cell envelope functions. The EMBO Journal, 39(13):e104926, July 2020.

16. S. Zhu, C. J. Bradfield, A. Maminska, E.-S. Park, B.-H. Kim, P. Kumar, S. Huang, M. Kim, Y. Zhang, J. Bewersdorf, et al. Native architecture of a human GBP1 defense complex for cell-autonomous immunity to infection. Science, 383(6686):eabm9903, March 2024.

17. J. C. Santos, M. S. Dick, B. Lagrange, D. Degrandi, K. Pfeffer, M. Yamamoto, E. Meunier, P. Pelczar, T. Henry, et al. LPS targets host guanylate-binding proteins to the bacterial outer membrane for non-canonical inflammasome activation. The EMBO Journal, 37(6): EMBJ201798089, February 2018.

18. M. S. Dickinson, M. Kutsch, L. Sistemich, D. Hernandez, A. S. Piro, D. Needham, C. F. Lesser, C. Herrmann, et al. LPS-aggregating proteins GBP1 and GBP2 are each sufficient to enhance caspase-4 activation both in cellulo and in vitro. Proceedings of the National Academy of Sciences, 120(15):e2216028120, April 2023.

19. T. Kuhm, C. Taisne, C. de Agrela Pinto, L. Gross, E. A. Giannopoulou, S. T. Huber, E. Pardon, J. Steyaert, S. J. Tans, et al. Structural basis of antimicrobial membrane coat assembly by human GBP1. Nature Structural & Molecular Biology, 32(1):172–184, January 2025.

20. J. Began, A. Boegli, L. Anton, D. Biriukov, J. Vairamuthu, J. Carlos Santos, et al. Caspase-4 binds to lps membranes with positive curvature for non-canonical inflammasome activation. Immunity, 59(3):542–558.e8, 2026.

21. M. Kutsch, S. Ince and C. Herrmann. Homo and hetero dimerisation of the human guanylate-binding proteins hGBP-1 and hGBP-5 characterised by affinities and kinetics. The FEBS Journal, 285(11):2019–2036, 2018.

22. B. Prakash, G. J. K. Praefcke, L. Renault, A. Wittinghofer and C. Herrmann. Structure of human guanylate-binding protein 1 representing a unique class of GTP-binding proteins. Nature, 403(6769):567–571, February 2000.

23. B. Prakash, L. Renault, G. J. Praefcke, C. Herrmann and A. Wittinghofer. Triphosphate structure of guanylate-binding protein 1 and implications for nucleotide binding and GTPase mechanism. The EMBO Journal, 19(17):4555–4564, September 2000.

24. A. Ghosh, G. J. K. Praefcke, L. Renault, A. Wittinghofer and C. Herrmann. How guanylate-binding proteins achieve assembly-stimulated processive cleavage of GTP to GMP. Nature, 440(7080):101–104, March 2006.

25. M. Weismehl, X. Chu, M. Kutsch, P. Lauterjung, C. Herrmann, M. Kudryashev, et al. Structural insights into the activation mechanism of antimicrobial GBP1. The EMBO Journal, 43(4):615–636, January 2024.

26. S. Shydlovskyi, A. Y. Zienert, S. Ince, C. Dovengerds, A. Hohendahl, J. M. Dargazanli, Blum, S. D. Günther, N. Kladt, M. Stürzl, A. C. Schauss, M. Kutsch, A. Roux, G. J. K. Praefcke, et al. Nucleotide-dependent farnesyl switch orchestrates polymerization and membrane binding of human guanylate-binding protein 1. Proceedings of the National Academy of Sciences, 114(28):E5559–E5568, July 2017.

27. C. Lorenz, S. Ince, T. Zhang, A. Cousin, R. Batra-Safferling, L. Nagel-Steger, C. Herrmann, et al. Farnesylation of human guanylate-binding protein 1 as safety mechanism preventing structural rearrangements and uninduced dimerization. The FEBS Journal, 287(3):496–514, 2020.

28. S. V. Valeva, M. Degabriel, F. Michal, G. Gay, J. R. Rohde, F. Randow, B. Lagrange, et al. Comparative study of GBP recruitment on two cytosol-dwelling pathogens, Francisella novicida and Shigella flexneri highlights differences in GBP repertoire and in GBP1 motif requirements. Pathogens and Disease, 81:ftad005, January 2023.

29. C. Hamers-Casterman, T. Atarhouch, S. Muyldermans, G. Robinson, C. Hammers, E. B. Songa, N. Bendahman, et al. Naturally occurring antibodies devoid of light chains. Nature, 363(6428):446–448, June 1993.

30. S. Muyldermans. Nanobodies: Natural Single-Domain Antibodies. Annual Review of Biochemistry, 82(1):775–797, June 2013.

31. U. Rothbauer, K. Zolghadr, S. Tillib, D. Nowak, L. Schermelleh, A. Gahl, N. Backmann, K. Conrath, S. Muyldermans, M. C. Cardoso, et al. Targeting and tracing antigens in live cells with fluorescent nanobodies. Nature Methods, 3(11):887–889, November 2006.

32. T. Pleiner, M. Bates, S. Trakhanov, C.-T. Lee, J. E. Schliep, H. Chug, M. Böhning, H. Stark, H. Urlaub, et al. Nanobodies: site-specific labeling for super-resolution imaging, rapid epitope-mapping and native protein complex isolation. eLife, 4:e11349, December 2015.

33. H. Götzke, M. Kilisch, M. Martínez-Carranza, S. Sograte-Idrissi, A. Rajavel, T. Schlichthaerle, N. Engels, R. Jungmann, P. Stenmark, F. Opazo, et al. The ALFA-tag is a highly versatile tool for nanobody-based bioscience applications. Nature Communications, 10(1):4403, September 2019.

34. S. G. F. Rasmussen, H.-J. Choi, J. J. Fung, E. Pardon, P. Casarosa, P. S. Chae, B. T. DeVree, D. M. Rosenbaum, F. S. Thian, T. S. Kobilka, A. Schnapp, I. Konetzki, R. K. Sunahara, S. H. Gellman, A. Pautsch, J. Steyaert, W. I. Weis, et al. Structure of a nanobody-stabilized active state of the β2 adrenoceptor. Nature, 469(7329):175–180, January 2011.

35. T. Uchański, E. Pardon and J. Steyaert. Nanobodies to study protein conformational states. Current Opinion in Structural Biology, 60:117–123, February 2020.

36. A. A. Gil, C. Carrasco-López, L. Zhu, E. M. Zhao, P. T. Ravindran, M. Z. Wilson, A. G. Goglia, J. L. Avalos, et al. Optogenetic control of protein binding using light-switchable nanobodies. Nature Communications, 11(1):4044, August 2020.

37. T. Uchański, S. Masiulis, B. Fischer, V. Kalichuk, U. López-Sánchez, E. Zarkadas, M. Weckener, A. Sente, P. Ward, A. Wohlkönig, T. Zögg, H. Remaut, J. H. Naismith, H. Nury, W. Vranken, A. R. Aricescu, E. Pardon, et al. Megabodies expand the nanobody toolkit for protein structure determination by single-particle cryo-EM. Nature Methods, 18(1):60–68, January 2021.

38. W. Cui, E. Braun, W. Wang, J. Tang, Y. Zheng, B. Slater, N. Li, C. Chen, Q. Liu, B. Wang, X. Li, Y. Duan, Y. Xiao, R. Ti, D. Hotter, X. Ji, L. Zhang, J. Cui, Y. Xiong, D. Sauter, Z. Wang, F. Kirchhoff, et al. Structural basis for GTP-induced dimerization and antiviral function of guanylate-binding proteins. Proceedings of the National Academy of Sciences, 118(15): e2022269118, April 2021.

39. T. I. Kuhm. Double trouble: Structural basis of Guanylate Binding Proteins targeting pathogen membranes. 2024.

40. L. Sistemich, L. Dimitrov Stanchev, M. Kutsch, A. Roux, T. Günther Pomorski and C. Herrmann. Structural requirements for membrane binding of human guanylate-binding protein 1. The FEBS Journal, 288(13):4098–4114, 2021.

41. E. Pardon, T. Laeremans, S. Triest, S. G. F. Rasmussen, A. Wohlkönig, A. Ruf, S. Muyldermans, W. G. J. Hol, B. K. Kobilka, et al. A general protocol for the generation of Nanobodies for structural biology. Nature Protocols, 9(3):674–693, March 2014.

42. M.-P. Lefranc, C. Pommié, M. Ruiz, V. Giudicelli, E. Foulquier, L. Truong, V. Thouvenin-Contet, et al. IMGT unique numbering for immunoglobulin and T cell receptor variable domains and Ig superfamily V-like domains. Developmental and Comparative Immunology, 27(1):55–77, January 2003.

43. S. D. Krämer, J. Wöhrle, C. Rath and G. Roth. Anabel: An Online Tool for the Real-Time Kinetic Analysis of Binding Events. Bioinformatics and Biology Insights, 13: 1177932218821383, January 2019.

44. A. Bharadwaj, R. d. Bruin and A. J. Jakobi. Confidence-guided cryo-EM map optimisation with LocScale-2.0. bioRxiv, page 2025.09.11.674726, September 2025.

45. E. C. Meng, T. D. Goddard, E. F. Pettersen, G. S. Couch, Z. J. Pearson, J. H. Morris, et al. UCSF ChimeraX: Tools for structure building and analysis. Protein Science, 32(11):e4792, 2023.

46. J. Jumper, R. Evans, A. Pritzel, T. Green, M. Figurnov, O. Ronneberger, K. Tunyasuvunakool, R. Bates, A. Žídek, A. Potapenko, A. Bridgland, C. Meyer, S. A. A. Kohl, A. J. Ballard, Cowie, B. Romera-Paredes, S. Nikolov, R. Jain, J. Adler, T. Back, S. Petersen, D. Reiman, E. Clancy, M. Zielinski, M. Steinegger, M. Pacholska, T. Berghammer, S. Bodenstein, D. Silver, O. Vinyals, A. W. Senior, K. Kavukcuoglu, P. Kohli, et al. Highly accurate protein structure prediction with AlphaFold. Nature, 596(7873):583–589, August 2021.

47. M. Mirdita, K. Schütze, Y. Moriwaki, L. Heo, S. Ovchinnikov and M. Steinegger. ColabFold: making protein folding accessible to all. Nature Methods, 19(6):679–682, June 2022.

48. R. J. Read, C. Millán, A. J. McCoy and T. C. Terwilliger. Likelihood-based signal and noise analysis for docking of models into cryo-EM maps. Acta Crystallographica Section D Structural Biology, 79(4):271–280, April 2023.

49. A. Fadini, M. Li, A. J. McCoy, S. Banjara, H. Okumura, E. Napier, P. Fontana, A. R. Khan, L. Jovine, T. C. Terwilliger, R. J. Read, D. R. Hekstra, et al. AlphaFold as a prior: experimental structure determination conditioned on a pretrained neural network. Nature Methods, 23(4):785–795, April 2026.

50. P. Emsley, B. Lohkamp, W. G. Scott and K. Cowtan. Features and development of Coot. Acta Crystallographica. Section D, Biological Crystallography, 66(Pt 4):486–501, April 2010.

51. G. Chojnowski. gapTrick—structural characterization of protein–protein interactions using AlphaFold. Bioinformatics, 41(9):btaf532, September 2025.

52. K. Yamashita, C. M. Palmer, T. Burnley and G. N. Murshudov. Cryo-EM single-particle structure refinement and map calculation using Servalcat. Acta Crystallographica Section D: Structural Biology, 77(10):1282–1291, October 2021.

53. A. Dümmler, A.-M. Lawrence and A. de Marco. Simplified screening for the detection of soluble fusion constructs expressed in E. coli using a modular set of vectors. Microbial Cell Factories, 4:34, December 2005.

54. E. R. Geertsma and R. Dutzler. A Versatile and Efficient High-Throughput Cloning Tool for Structural Biology. Biochemistry, 50(15):3272–3278, April 2011.

55. R. Zell and H. J. Fritz. DNA mismatch-repair in Escherichia coli counteracting the hydrolytic deamination of 5-methyl-cytosine residues. The EMBO Journal, 6(6):1809–1815, June 1987.

